# Modular gene tagging in *C. elegans*

**DOI:** 10.64898/2026.01.15.699741

**Authors:** Adam Hefel, Kevin Kruse, Kaden Wall, Soren B. Jorgensen, Kam Hoe Ng, Ryan Stolley, Matthew S. Rich, Erik M. Jorgensen

**Affiliations:** School of Biological Sciences, University of Utah, Salt Lake City, Utah, USA; Howard Hughes Medical Institute, University of Utah, Salt Lake City, Utah, USA; Department of Chemistry, University of Utah, Salt Lake City, Utah, USA

**Keywords:** *C. elegans* genome engineering, gene tagging, PhIT, PhiC31, Dre, B3 recombinase

## Abstract

Tagging a gene endogenously can identify when the gene is expressed and where the protein is localized. CRISPR is the primary tool for generating tags of endogenous genes, but it is error-prone and requires unique reagents for each gene and tag. Recombinases can insert DNA in an error-free and modular manner. Here, we tested eight recombinases for germline function in the nematode *C. elegans*, and introduce PhIT, a recombinase-based method for protein tagging. First, a short 39bp PhiC31 *attB* landing pad is inserted into the locus by CRISPR. This strain is a resource which can be used to insert a variety of modular tags. Second, tags are inserted by the integrase PhiC31, and in tandem, extraneous backbone sequences are removed by a tyrosine recombinase. Current modular tags include seven different fluorescent proteins, FLP-regulated cell-specific expression constructs, and degron tags. Importantly, tags can be inserted by genetic crosses instead of by microinjection.

## Introduction

Three protein analyses are typically used to characterize function: mutant phenotype, protein interaction, and cellular localization. High-throughput methods have been developed to generate gene knockouts and identify binding partners (Howson et al., 2005; Huh et al., 2003). Although methods for protein localization have advanced, there are formidable barriers. For fixed cells, antibody staining is the preferred method for visualization, but production of antibodies requires considerable costs and validation. An alternative is to introduce a transgene tagged with a fluorescent protein which can localize proteins in living samples. However, transgenes are not embedded in the normal chromatin context and are often expressed under the control of heterologous promoters.

It is now possible to tag endogenous genes using CRISPR (Cong et al., 2013; Doudna and Charpentier, 2014; Mali et al., 2013). However, current CRISPR approaches are often restricted to one-off experiments for tagging a single gene. First, double-strand break repair by synthesis-dependent strand annealing is non-processive and error-prone, which limits insertion size (Frøkjaer-Jensen et al., 2008; Paix et al., 2017). Second, significant time and resources are required to construct and introduce editing reagents into the germline. Third, introduction of different tags into a single gene requires independent rounds of CRISPR. Editing of many genes and with diverse tags is not yet practical.

By contrast, recombinases insert DNA into the genome with near-perfect fidelity, and have become workhorses for genome rearrangements in mouse (Danielian et al., 1998), *Drosophila* (Golic and Lindquist, 1989; Nern et al., 2011) and roundworm (Davis et al., 2008; Hoier et al., 2000; Nonet, 2023, 2020; Voutev and Hubbard, 2008; Yang et al., 2022). There are two major families of recombinases: tyrosine recombinases and serine integrases, named for the residue at the enzymatic site. Tyrosine recombinases act at target sites composed of inverted repeats and an intervening ‘overlap’ sequence that provides specificity and orients the site (Sauer, 1998). Matching recombinase target sequences can be used to promote excision (Cre/*lox*, Sauer, 1998), insertion (FLP RMCE, Nonet, 2020), or DNA rearrangements (FLP-on, Davis et al., 2008). These rearrangements are reversible because tyrosine recombinases regenerate the target site, which can be used again for recombination.

In contrast to tyrosine recombinases for which recombination is reversible, serine integrases, such as PhiC31 and Bxb1, are not reversible. Integrases mediate recombination between two different target sites, the bacterial and phage attachment sites (*attB* and *attP*). Recombination between these sites alters the original sites, generating *attL* and *attR* sites, which cannot be reversibly recombined without additional factors (Li et al., 2018). Thus, the insertion is fixed in the genome. The benefit of using recombinases for modifying the genome is that they do not rely on DNA damage repair, and are therefore error-free. Likewise, this method is not subject to the same limitations on insert size as CRISPR (Paix et al., 2016).

Here, we tested eight recombinases to identify those that work well in the germline of *C. elegans*, including five previously untested recombinases (Dre, Vika, KD, B3 and Bxb1) (Abel and Theologis, 1996; Chen et al., 1986; Karimova et al., 2013; Keravala et al., 2006; Nern et al., 2011; Sauer and McDermott, 2004). These recombinases were then used to insert tags into the genome. This method, called PhIT (PhiC31-mediated Integration of Transgenes/Tags), uses a small *attB* tag (39bp, previously shown to have no native or pseudo-sites in *C. elegans* (Nonet, 2024)) generated by CRISPR in the gene of interest(Ghanta and Mello, 2020; Paix et al., 2017). The *attB* tag functions as a landing pad for insertion of a variety of tags into the gene of interest with minimal scars. Plasmid DNA with tag sequences are injected into *C. elegans,* where it is assembled into extrachromosomal arrays and transmitted as a mini-chromosome during cell division. In PhIT, a tyrosine recombinase excises the tag from the extrachromosomal array, and a serine integrase inserts the tag of interest into the gene. A large selection of plasmid reagents encoding a variety of tags (including fluorescent proteins, degrons, or cell-specific expression constructs) can be inserted as interchangeable modules at the N- or C-termini of proteins.

Importantly, this method makes genome engineering in *C. elegans* broadly accessible, since we developed strains that enable *C. elegans* tags to be inserted into genes by simple genetic crosses. The methods described here could be adapted to other model organisms to facilitate genome engineering applications more broadly in the scientific and educational communities.

## Results

### Recombinase-mediated excision

We tested eight recombinases in the *C. elegans* germline: six tyrosine recombinases (B3, Cre, Dre, FLP, KD, and Vika), and two serine integrases (Bxb1, and PhiC31). DNA constructs were built to contain target sites for each of these recombinases flanking a marker, and each was integrated as a single copy in the genome. The transgene marker encodes a mutant collagen that causes a dominant ‘rolling’ phenotype (Rol) (Figure 1A) (Kramer and Johnson, 1993). Each of these strains was injected with plasmids expressing the cognate recombinase in the germline. When the recombinases bind the target sites, recombination between the aligned sites excises the Rol marker and generates non-rolling progeny (non-Rol, that appears ‘wild type’).

**Figure 1:**
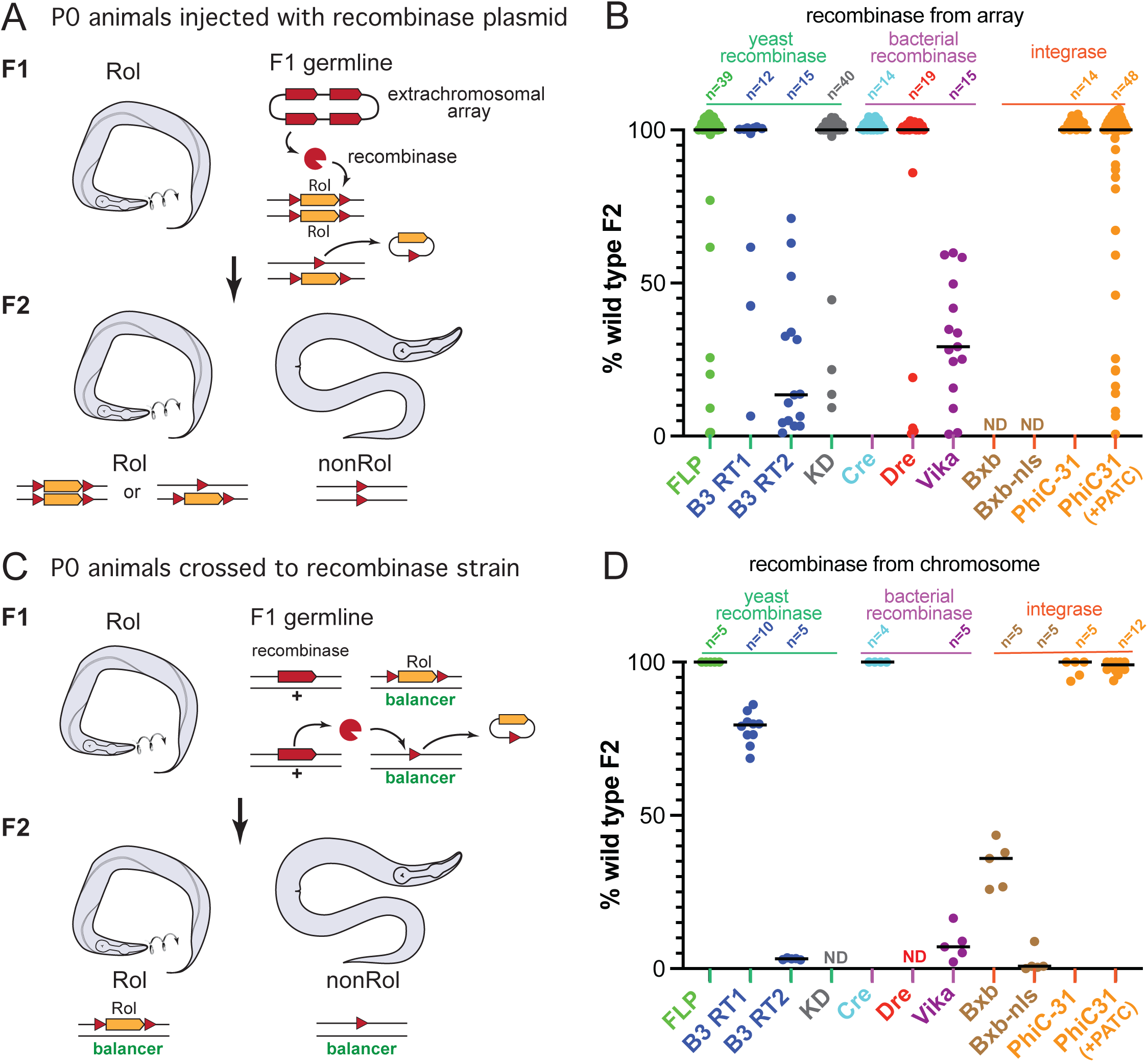
Recombinase excision assays. A transgene causing a dominant rolling (Rol) phenotype (*sqt–1(e1350)*) was inserted as a single-copy into chromosome I. The Rol markers were flanked by recombination targets for each of the recombinases. (A) Recombinase from multicopy array. Rol animals were injected with plasmids expressing the recombinases under a germline promoter (P*mex-5*). Recombination and excision of the Rol marker occurs in the F1 germline. Homozygous F2 progeny with excised transgenes move normally (non-Rol). (B) Excision rates of recombinases. Array-positive F1 animals were singled and assayed for fraction of non-Rol animals among the F2. Each point represents the proportion of non-Rol progeny from a single F1. Only F1s with some non-Rol progeny were scored to ensure the recombinase was present in the array. (C) Recombinase from a single-copy transgene. Males expressing the recombinase from a single copy of an integrated transgene were crossed to Rol hermaphrodites. Excision occurs in the F1 germline. Balanced F2 worms were scored for the Rol phenotype. (D) Excision rates from F1s with a single copy of the excision marker. Each point represents the proportion of non-Rol animals among balanced F2s from a single F1. n-values represent number of singled F1s, lines indicate median values. ND, no data.

DNA injected into *C. elegans* parental animals (P0) forms a circular DNA molecule in the germline, and is incorporated as a minichromosome in the zygote of the progeny (F1 generation). Because the recombinase is only expressed in the germline, these F1 animals are almost all Rol, with somatic markers still intact. If the recombinase excises the Rol transgene in germline cells that will become sperm and the oocytes, non-Rol progeny will be observed among the F2 animals.

The excision rate for each recombinase was calculated as the percent of non-Rol F2 progeny from singled F1s (Figure 1B). The recombinases FLP and Cre are commonly used for transgene excision or cell-specific expression in *C. elegans*; as expected, these recombinases generated high proportions of non-Rol F2 progeny (100% and 84%, respectively). B3, Dre, and KD, also excised the transgene efficiently (83%, 80%, and 92%, respectively), whereas Vika produced few non-Rol animals (31%). The serine integrase PhiC31 generated 100% non-Rol progeny. A PhiC31 construct optimized for germline expression (intron containing periodic AT clusters (PATC), (Frøkjær-Jensen et al., 2016)) did not improve recombination frequency (81%).

In rare cases, F1 animals were non-Rol (8.8%, 19/216). These non-Rol F1 did not always breed true (11%, 2/19) suggesting that the recombinase was expressed early in the P0 or zygote, excising the transgene in the somatic lineages but not in the F1 germline. Almost half of the non-Rol F1 were generated by KD recombinase (42%, 8/19). KD was one of the most efficient of the tyrosine recombinases in this study (Figure 1B) and it is possible that it excised the Rol marker early because of its high activity.

There are two drawbacks to this assay. First, extrachromosomal arrays can carry 100’s of gene copies, so recombinase expression from arrays may vary. Second, the rate of excision cannot be determined for a single copy of the reporter: because the Rol marker is dominant, generation of non-Rol F2 requires two excision events (one for each chromosome).

To test the efficiency of single-copy recombinase expression constructs, the recombinase expression constructs were integrated into the genome. Constructs for Dre and KD could not be integrated, possibly due to lethality when incorporated into the genome. The recombinase-expressing constructs were crossed into the Rol excision marker strains. All F1 progeny were Rol, suggesting that excision does not occur in the F1 zygote, but rather the F1 germline.

Excision rate was assayed as the percent of non-Rol F2 progeny from each singled F1 (Figure 1D). These results confirm the recombinase efficiencies from plasmid injection: Cre and FLP excised the Rol marker in 100% of the progeny. PhiC31 excised the Rol marker in 98% of progeny. B3, Vika, and Bxb1 exhibited lower rates of excision (B3-*B3RT1* 78%, B3-*B3RT2* 3%, Vika 8%, Bxb1 34%, Bxb1-NLS 2%). These data indicate the recombinase enzymes function in the germline when expressed from multicopy arrays or as single integrated expression constructs.

### Single-copy transgene insertions

Recombinases were tested as reagents for generating single-copy transgene insertions. PhIT (PhiC31-mediated Integration of Transgenes / Tags) uses two recombinases: a PhiC31 integrase to insert the transgene, and a tyrosine recombinase to remove plasmid backbone sequences.

Serine integrases, like PhiC31, promote unidirectional recombination between *attP* and *attB* target sites. PhiC31 was the most efficient integrase in our excision assays (comparing PhiC31, Bxb1 and Bxb1-nls in Figure 1D). We generated three minimal PhiC31 *attB* landing pads (38 bp) by CRISPR on chromosomes I, II, and IV at positions previously characterized as permissive for gene expression (Figure S1; Table S1 for insertion site sequences (Frøkjaer-Jensen et al., 2008)). These strains also carry mutations in the *unc-119* gene as a rescuable marker. Transgene constructs carry an *unc-119(+)* rescuing construct to facilitate insertion selection. These were generated by SapTrap DNA assembly (Davis and Jorgensen, 2025; Schwartz and Jorgensen, 2016), in which the donor plasmids can be mixed and matched as modules to assemble a variety of transgene constructs (Figure S2). To assay insertion efficiency by PhIT, we first assembled constructs expressing the mKate2 fluorescent protein under the control of a dopamine neuron promoter (P*dat–1::mKate2*).

Tyrosine recombinases were compared for their ability to promote insertions by PhiC31. Bacterial DNA could alter gene expression and should not be incorporated with the gene tag. Excising the backbone has the added benefit of reducing the extrachromosomal array to ‘unit circles’. These ‘unit circles’ act as freely diffusing substrates for integration (Figure 2B). The mKate transgene was flanked by recombination targets for three different recombinases, KD, Dre, and B3 (*KDRT*, *rox*, and *B3RT1* respectively). Vika was not tested since it performed poorly in the excision assays.

**Figure 2:**
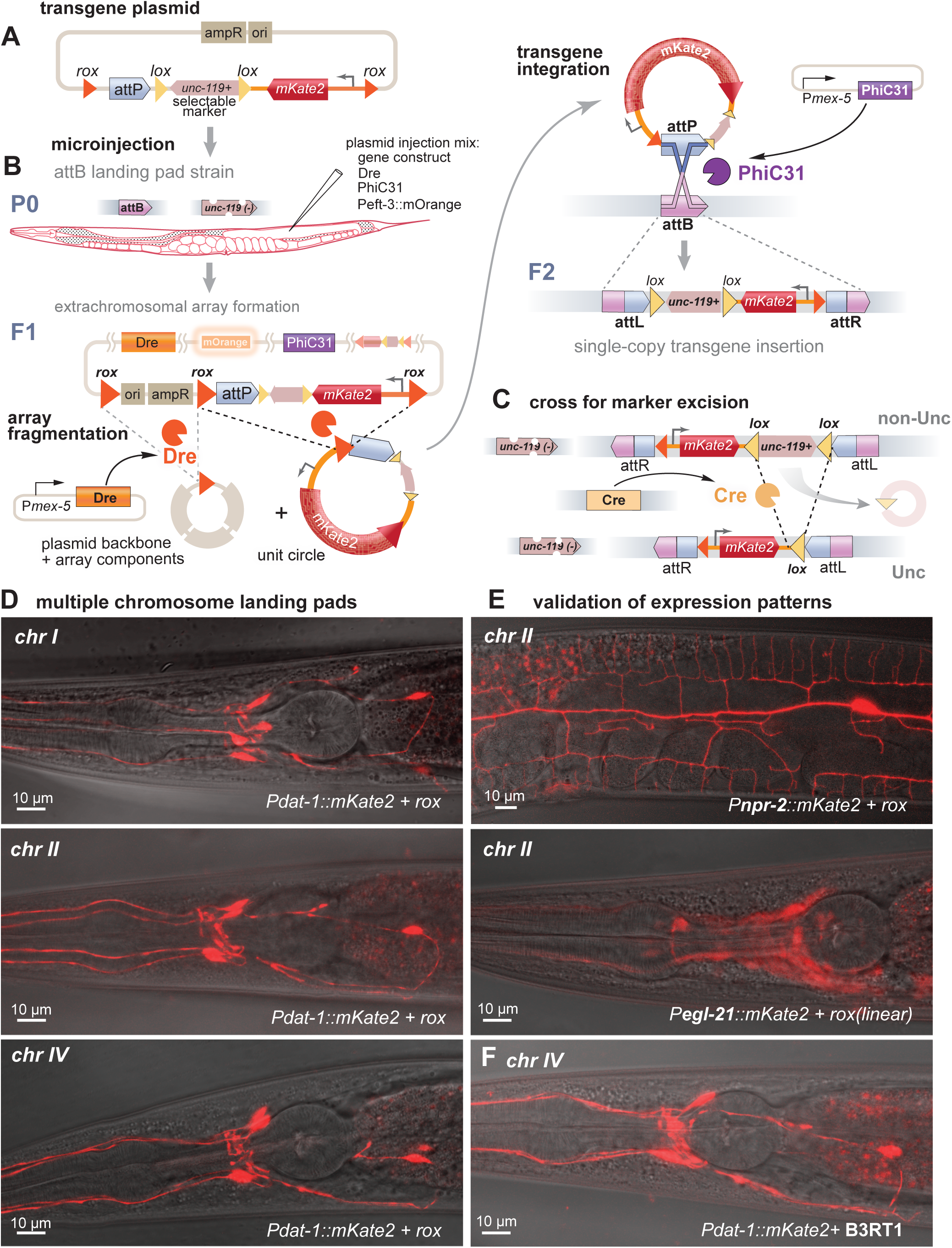
Single-copy transgene insertions. (A) Transgene. The plasmid contains the fluorescent transgene, a *Cb-unc–119(+)* selection marker, and an *attP* recombination target. This construct was flanked by tyrosine recombinase targets for KD, B3, or Dre. *rox* sites for the Dre recombinase are shown as an example. (B) Microinjection. Animals carrying the *attB* landing pads and an *unc–119(ox819)* mutation were injected with plasmids encoding PhiC31 integrase, a tyrosine recombinase (Dre, KD, or B3), and an orange fluorescent array marker (pWD540). DNA injected into the gonad of the worm forms extrachromosomal arrays. These arrays are fragmented by the tyrosine recombinase in the F1 germline to form ‘unit circles’ containing the transgene but lacking the plasmid backbone. Fragmentation also promotes diffusion of the minicircle versions of the construct and reduces and destabilizes the extrachromosomal array. ‘Unit circles’ containing the *attP* site are integrated into the genome by PhiC31-mediated recombination of the *attP* with the *attB* site in the genome. Successful integration will generate non-Unc F2 animals, which will also lack the orange fluorescent array marker. (C) Excision of selection marker. The *C. briggsae unc-119(+)* gene is flanked by *lox* sites and can be removed by crossing in Cre recombinase. (D) Landing pads. Examples of P*dat-1::mKate2* insertions at *attB* insertion site on different chromosomes, generated with PhiC31 and Dre recombinase. (E) Expression patterns. The insertion sites are permissive for specific expression of mKate2 (inserted by PhiC31 and Dre recombinase): P*npr-2* expresses in PVD neurons, P*egl-21* is ubiquitously expressed in neurons, and can be seen here in cell bodies of the anterior ganglia. (F) Tyrosine recombinase targets. Use of the B3 recombinase target site *B3RT1* in the construct generates and identical expression pattern as use of the Dre target site *rox* (panel D bottom) for mKate expression in dopamine neurons (P*dat-1::mKate2*).

Plasmids encoding the fluorescent transgene, PhiC31, the tyrosine recombinase, and an array marker, were injected into the gonads of a strain containing an *attB* landing pad (Figure 2B). Array-bearing F1 progeny were identified by *unc-119* rescue. F2 progeny with an integrated transgene were identified by *unc-119* rescue and absence of the array marker. Insertion was confirmed by PCR and the strains were crossed to Cre-expressing animals to remove the *unc–119* selection (Figure 2C).

Insertion rates were calculated as the percent of F1 arrays that produced tagged F2 progeny. Dre and B3 had similar insertion rates (9.1% and 9.8%, respectively). Injection of linearized fragments instead of circular plasmids was more successful (Dre 27.3%) and produced an identical expression pattern (Figure 2E and S3). In contrast to Dre and B3, KD produced insertions at a much lower rate (1.5%) (Table 1). This was surprising, since KD exhibited a high rate of excision in our previous assay (Figure 1A). It is possible that KD is too efficient; that is, it resolves the array into small circular DNA molecules so rapidly that they are lost by cell division before they can be integrated.

**Table 1.** mKate2 transgene Insertion.

| RT/recombinase | # injected animals | # WT F1 animals | # WT F1 w/ insertion | percent WT F1 insertion |
| --- | --- | --- | --- | --- |
| B3RT2/B3 | 175 | 80 | 9 | **9.75%** |
| Rox/Dre | 140 | 197 | 18 | **9.13%** |
| KDRT/KD | 150 | 146 | 2 | **1.46%** |

Expression was evaluated at each *attB* insertion site to validate these reagents (Figure 2D).

Insertions at all three landing pads exhibited specific expression in dopamine neurons (P*dat–1::mKate2*) (Figure 2D and Figure S3). Expected expression patterns were observed for transgenes regulated by different promoters: expressed broadly in all neurons (P*egl–21*) and narrowly in the highly ramified PVD neuron (P*npr–2*) (Figure 2E; Figure S3), indicating that the chromatin context surrounding these insertion sites does not disrupt promoter specificity.

Additionally, recombinase target sites did not appear to alter expression, since Dre and B3 generated insertions with identical expression patterns (Figure 2D bottom panel, Figure 2F).

### Tagging genes by injection

In addition to insertion of transgenes, PhIT is useful for tagging endogenous genes: First, insertion is mediated by recombination, rather than by error-prone double-strand break repair. Second, bacterial DNA from the plasmid, which could alter gene expression or interfere with translation, is removed by the recombinase. Third, it is modular – the landing pad in the gene can be used to insert a wide variety of gene tags.

We developed PhIT reagents for N-terminal and C-terminal tagging by testing tags in synaptobrevin (SNB-1) and a V-ATPase subunit (UNC–32), respectively. These are synaptic vesicle proteins and have a distinctive subcellular localization, and the loci have been extensively characterized for gene tagging by CRISPR (Schwartz and Jorgensen, 2016). *attB* landing pads were made with CRISPR in an *unc-119(ox819)* background. To maintain the open reading frame, the PhiC31 *attB* target site was extended to 39bp (Figure S4). These *attB* sites were inserted after the native start codon of *snb–1*, and before the native stop codon of *unc–32* (Figure 3A and 3D).

**Figure 3:**
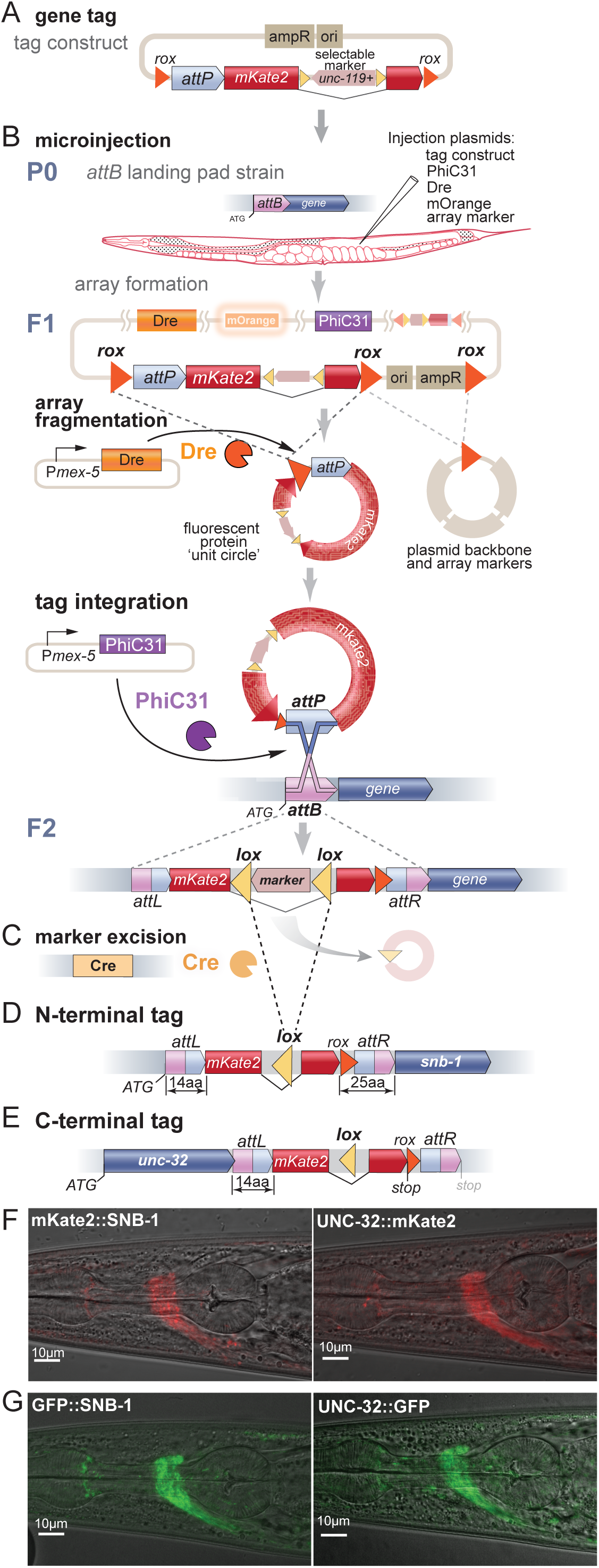
Gene tagging with PhIT. (A) Tagging construct. Example of an N-terminal tag using PhIT. The plasmid contains an *attP* recombination target in front of the fluorescent tag, and the *unc–119(+)* selection marker is inserted into an intron of the tag in the reverse orientation. This construct was flanked by the tyrosine recombinase target sites *rox* or *B3RT2*, for Dre and B3, respectively (*rox* sites shown in example). (B) Microinjection. Animals carrying the *attB* landing pads and an *unc-119(ox819)* mutation were injected with plasmids encoding PhiC31 integrase, a tyrosine recombinase (Dre or B3), and an orange fluorescent array marker. In the F1 germline, recombination of the *rox* sites causes the array to fragment and to generate ‘unit circles’ encoding just the fluorescent tag. PhiC31 recombines the *attB* and *attP* sites, leading to insertion of the tag into the gene. (C) Excision of selection marker. The *unc-119(+)* gene is flanked by *lox* sites and is removed by crossing or injecting Cre recombinase. (D) An example of N-terminal tag of SNB-1 using Dre recombinase. The *attL* recombination product is translated and encodes 14 aa sequence in front of the tag. The *rox* site and the *attR* recombination product are translated as a 25 aa linker between the tag and the protein. (E) An example of a C-terminal tag of UNC-32 using Dre recombinase. The *attL* recombination product encodes a 14 aa linker between the protein and the tag. The *rox* site and the *attR* recombination product are not translated because a stop site is encoded at the end of the tag. (F) Fluorescence images show nerve-ring localization of mKate2 for the SNB-1 N-terminal tag (left) and for the UNC-32 C-terminal tag (right). (G) Fluorescence images show nerve-ring localization of GFP for the SNB-1 N-terminal tag (left) and for the UNC-32 C-terminal tag (right).

Recombination of the *attB* and *attP* sites creates *attL* and *attR* chimeric sequences. These post-recombination sites are incorporated into the tagged gene and therefore were optimized to serve as linkers (Figure S4 and S5). For N-terminal tags the tyrosine recombinase target is incorporated into the linker with *attR* (Figure 3A). C–terminal tag constructs include a stop codon after the tag to prevent translation of the *attR* and recombinase target scar (Figure 3E). Tagging plasmids were assembled using SapTrap to enable modular cloning of fluorescent proteins (Figure S3, (Schwartz and Jorgensen, 2016)).

As with transgene insertions, plasmids encoding an array marker, the tag, PhiC31, and a tyrosine recombinase were injected into the gonad of *unc–119* mutants carrying the *attB* landing pad (Figure 3B). In the F2 generation, integration of the tag was identified by *unc–119(+)* rescue and loss of the fluorescent array marker. After confirming insertion, the strains were crossed to Cre-expressing animals to remove the *unc-119* selection (Figure 3C). Fluorescence for N-terminally tagged mKate2::SNB-1 and C-terminally tagged UNC-32::mKate2 was located at synapses in the nerve ring, as expected (Figure 3F; Figure S3).

Insertion rates were calculated as the percent of array positive F1 animals that produced tagged genes. Only the tyrosine recombinases Dre and B3 were tested for gene tagging since KD exhibited low rates of transgene insertion. Dre was four-fold more efficient than B3 for array fragmentation (2.8% versus 10.8% stable arrays, respectively), and ten-fold more efficient at generating tags (5% versus 0.5%, respectively). The high rates of array fragmentation coupled to high insertion rate suggests that the number and mobility of the small DNA circles is essential for efficient gene insertion. Because of its high efficiency, Dre is recommended for tagging genes by injection.

### Modules

The power of PhIT tagging is its modularity: the attB-tagged gene can be used as a resource to drop-in different tags. We have constructed inserts that encode multiple colors, cell-specific tagging modules, and cell-specific degradation modules (Figure S6). To determine whether proteins colocalize, requires them to be tagged with different fluorophores. We generated SapTrap modules for a variety of different fluorescent proteins (Figure S2). Here, for example, the same strains harboring *attB* landing pads in *unc-32* and *snb-1* were reused to insert GFP at the same sites (Figure 3G and Table 2).

**Table 2.**
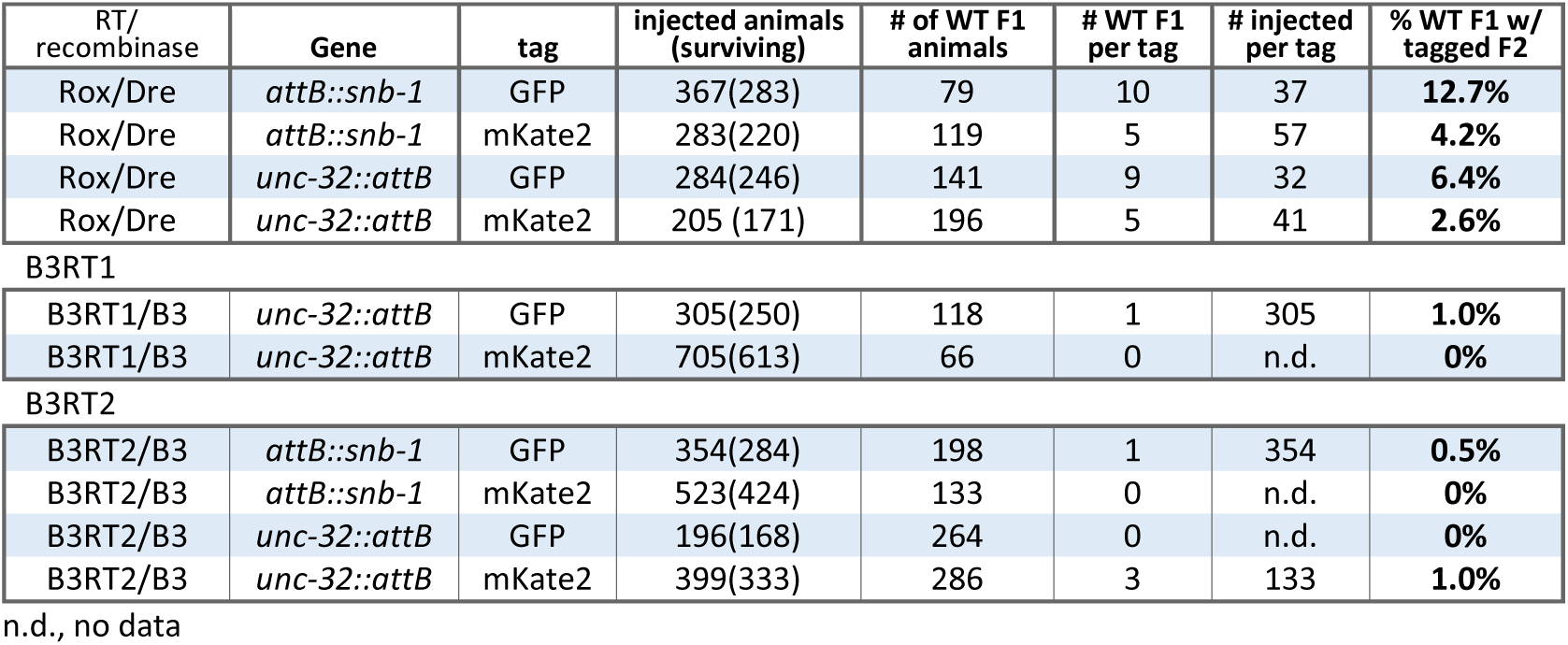
mKate2 and GFP tag insertion by microinjection.

| RT/recombinase | Gene | tag | injected animals (surviving) | # of WT F1 animals | # WT F1 per tag | # injected per tag | % WT F1 w/ tagged F2 |
| --- | --- | --- | --- | --- | --- | --- | --- |
| Rox/Dre | *attB::snb-1* | GFP | 367(283) | 79 | 10 | 37 | **12.7%** |
| Rox/Dre | *attB::snb-1* | mKate2 | 283(220) | 119 | 5 | 57 | **4.2%** |
| Rox/Dre | *unc-32::attB* | GFP | 284(246) | 141 | 9 | 32 | **6.4%** |
| Rox/Dre | *unc-32::attB* | mKate2 | 205 (171) | 196 | 5 | 41 | **2.6%** |
| B3RT1 |  |  |  |  |  |  |  |
| B3RT1/B3 | *unc-32::attB* | GFP | 305(250) | 118 | 1 | 305 | **1.0%** |
| B3RT1/B3 | *unc-32::attB* | mKate2 | 705(613) | 66 | 0 | n.d. | **0%** |
| B3RT2 |  |  |  |  |  |  |  |
| B3RT2/B3 | *attB::snb-1* | GFP | 354(284) | 198 | 1 | 354 | **0.5%** |
| B3RT2/B3 | *attB::snb-1* | mKate2 | 523(424) | 133 | 0 | n.d. | **0%** |
| B3RT2/B3 | *unc-32::attB* | GFP | 196(168) | 264 | 0 | n.d. | **0%** |
| B3RT2/B3 | *unc-32::attB* | mKate2 | 399(333) | 286 | 3 | 133 | **1.0%** |

n.d., no data

Cell-specific tagging is particularly important for resolving the subcellular location of broadly expressed proteins. FLP-on constructs allow for tissue fluorescent tagging (Davis et al., 2008; Schwartz and Jorgensen, 2016). In these constructs, FLP recombinase targets (FRTs) flank a cassette that separates the fluorescent tag from the gene-of-interest. This arrangement prevents expression of the fluorescent tag. However, excision of the cassette by FLP in cells where it is expressed causes fusion of the fluorescent tag with the tagged gene.

For cell-specific tagging using N-terminal FLP-on constructs, the fluorescent protein is expressed from an operon at the 5’ end of the gene (Figure 4A). The fluorescent tag is fused to a PEST degron, causing it to be rapidly degraded (Li et al., 1998). When FLP is expressed, these intervening sequences are excised, leading to fusion of the tag with the gene-of-interest. The N-terminal FLP-on construct was used to tag SNB-1 with PhIT (*mKate-2::FLP-on::snb-1*), and FLP recombinase was expressed in either all neurons (P*snt-1::FLP*) or just GABA neurons (P*unc-47::FLP*) (Fragoso-Luna et al., 2023). As expected, mKate2-tagged SNB-1 was observed in the synapse-rich nerve ring of all neurons, or just the GABA neurons, respectively (Figure 4B).

**Figure 4:**
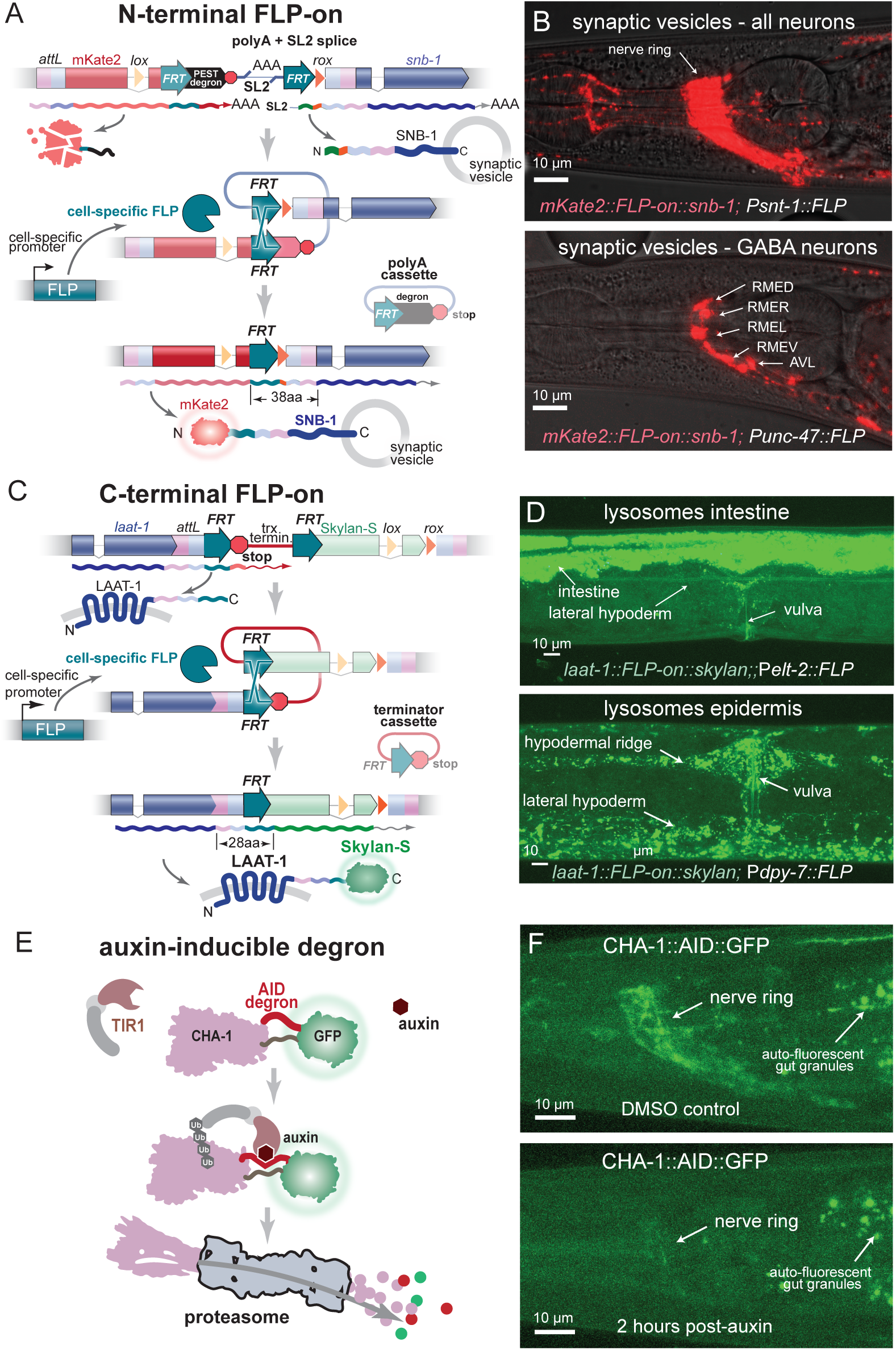
Modular tags: cell-specific tagging and cell-specific degradation. Cell-specific tagging modules using FLP-on cassettes. (A) Example of FLP-on cell-specific tagging at the N-terminus of the synaptic vesicle protein SNB–1. In most cells the untagged gene is expressed from an operon under the regulation of its own promoter using trans-spliced leader (SL2). The fluorescent protein is transcribed as a separate polyadenylated mRNA; however, a PEST degron at the C–terminus rapidly degrades the fluorescent protein. Tissue-specific expression of FLP recombinase rearranges the gene structure and excises the intervening sequence in those particular cells and joins the tag to the protein. Therefore, the tagged protein is only fluorescent in cells where the FLP transgene is expressed. (B) FLP–on tagging of SNB-1 with mKate. Confocal images of FLP-on::mKate2 tagged SNB-1 using FLP expressed in all neurons (top, P*snt–1::FLP*) or only in GABA neurons (bottom, P*unc–47::FLP*). (C) Example of FLP–on cell-specific tagging at the C-terminus of the lysosomal protein LAAT–1. C–terminal FLP–on tags use a ‘stop cassette’. FLP targets flank a stop codon and terminator at the C–terminus of the gene to prevent transcription of the tag. Cell–specific expression of FLP excises the terminator from the DNA and attaches the coding sequence to the tag. The tag is only translated in cells where FLP is expressed. (D) FLP–on tag of LAAT–1 with Skylan–S at the C–terminus. Confocal images of strains using FLP expressed in the intestine (P*elt–2*, top) or the epidermis (P*dpy–7*, bottom). Cell specific degradation module. (E) An auxin-inducible degron (AID::GFP) was inserted by a cross as an internal tag in in the gene for choline acetyltransferase CHA1. The modified *Arabidopsis* auxin-dependent ubiquitin ligase adaptor (TIR1(F79G)) was expressed in neurons (P*snt-1*). When auxin is applied, TIR1 ubiquitinates CHA–1, and the tagged protein is degraded by the proteosome. (F) CHA–1 degradation after auxin treatment. Young-adult worms were transferred to plates with auxin (5–phenyl auxin) or DMSO control. Control and treated worms were imaged 2 hours after treatment. GFP fluorescence was nearly abolished from the nerve ring in treated worms.

For C-terminal FLP-on tags, an FRT-flanked transcriptional stop cassette is inserted at the 3’ end of the gene before the tag. In cells in which FLP is expressed, the stop codon and terminator are excised, resulting in a fusion of the tag and the protein-of-interest (Figure 4C). Using PhIT, the C-terminus of LAAT-1 (a ubiquitously expressed lysosomal protein) was tagged with a FLP-on Skylan-S tag. FLP was expressed in the intestine (using P*elt-2*:*:FLP*) or in the epidermis (P*dpy-7::FLP*) and fluorescence was observed in intestine or epidermis, respectively (Figure 4D).

Drug induced degradation modules are particularly useful for determining the function of proteins that are pleiotropic, function at different time points, or are lethal as null mutants. Auxin-inducible degron tags (AID) can lead to selective degradation of protein in specific cells and at specific times (Nishimura et al., 2009; Zhang et al., 2015). Cell-specific expression of the auxin-inducible ubiquitin ligase TIR1 limits degradation to specific cells with the timing of degradation dependent on auxin application. (Abel and Theologis, 1996; Tan et al., 2007). Here, we used the TIR1(F79G) variant, which only responds to 5-phenyl-auxin and exhibits increased specificity and sensitivity (Hills-Muckey et al., 2022; Uchida et al., 2018). 5-phenyl-auxin is expensive and can be synthesized in the lab (Sural et al., 2024); here we developed a synthesis protocol so that it can be easily synthesized locally (Figure S7).

To test the AID tag as a PhIT module, the biosynthetic enzyme for acetylcholine (CHA-1) was tagged with AID::GFP. Null mutants in *cha-1* are lethal, and we were unable to tag the protein at N- or C-termini. However, an internal tagging site tolerated an *attB* landing pad, and insertion of an AID::GFP tagging module (Figure 4E). A construct expressing TIR1 in neurons (P*snt-1::TIR1*(F79G) IV) was crossed into the strain. Addition of 5-phenyl auxin resulted in reduction of CHA-1::GFP fluorescence within 2 hours (Figure 4F) and animals exhibited a coiling behavior typical of viable alleles. The F1 progeny phenocopied *cha-1* null mutants and died after hatching.

### Tagging genes by cross

Engineering the *C. elegans* genome requires specialized equipment and non-trivial microinjection skills that limit its utility as a model organism. To reduce this hurdle, we developed reagents to perform gene tagging by genetic crosses. PhiC31 integrase and B3 recombinase were expressed in the germline from integrated transgenes. B3 recombinase was used because Dre appears to be lethal when expressed in the genome (Figure 1B).

In short, we generated strains that each carry a fluorescent protein tag in an extrachromosomal array (Figure 5A). These strains were crossed to *attB* landing pad strains and then to a strain expressing the recombinases in the germline. F2 animals carry all three components; leading to array fragmentation and tag integration at the *attB* landing pad. Animals carrying an intact array are paralyzed by histamine because they express the *Drosophila* histamine-chloride ion channel (El Mouridi et al., 2021; Gisselmann et al., 2002; Pokala et al., 2014). Integrants are selected because they carry the rescuing marker *unc-119+* and lack all other array markers.

**Figure 5:**
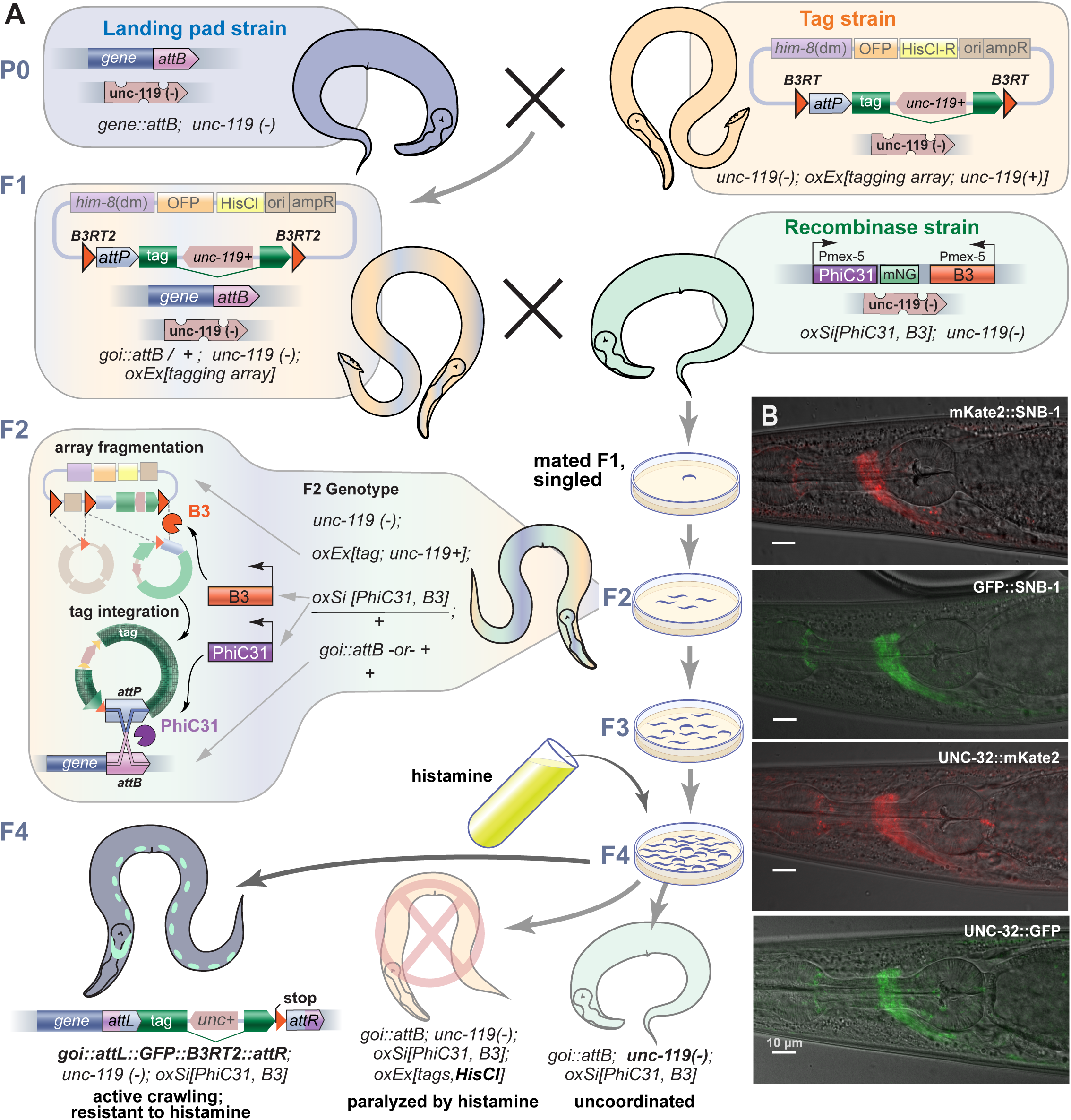
Tag insertion by cross. (A) C-terminal protein tagging using a PhiT cross. In the first cross, the *attB* tag landing pad strain, which is uncoordinated (*unc-119(-)*), is crossed to the tag array strain, which has an extra-chromosomal array consisting of the tag, a *him-8* piRNA sequence, and two array markers (OFP and HisCl). These animals have *unc-119(ox819)* in the background, which is rescued by the *unc-119(+)* in the intron of the tag to generate wild-type moving animals. The *him-8* piRNA knocks-down the *him-8* gene, which causes the strain to segregate males. The F1 males are mated with hermaphrodites expressing germline PhiC31 and B3. The array is fragmented by the tyrosine recombinase and the resulting unit circle containing the *attP* and the tag are integrated into the *attB* landing-pad. Plates are treated with histamine to paralyze animals carrying the extra-chromosomal array and active animals selected. Note that resistance to histamine is not a completely reliable marker for loss of the array, in some cases recombination by B3 eliminated the HisCl array marker (Table 3). Array loss was confirmed by absence of fluorescent markers. (B) Examples of tags made by crossing. UNC-32 and SNB-1 were tagged with mKate2 and GFP PhIT crosses, with expected fluorescence observed in the nerve ring of tagged animals.

**Table 3.** mKate2 and GFP insertion by crossing.

| Allele | Tag | Gene | mated F1s singled | plates w/ tagged progeny | % F1s w/ tagged progeny | plates w/ HisR progeny | HisR plates w/ no tags | % false positive HisR |
| --- | --- | --- | --- | --- | --- | --- | --- | --- |
| *oxEx2302* | N-term GFP | *attB::snb-1* | 25 | 6 | **24.0%** | 6 | 0 | 0% |
| *oxEx2300* | N-term mKate2 | *attB::snb-1* | 26 | 2 | **7.7%** | 16 | 14 | 88% |
| *oxEx2303* | C-term GFP | *unc-32::attB* | 23 | 10 | **43.5%** | 14 | 4 | 29% |
| *oxEx2301* | C-term mKate2 | *unc-32::attB* | 22 | 10 | **45.5%** | 18 | 8 | 44% |

Tags were inserted into the *snb-1* and *unc-32 attB* sites by two consecutive crosses to generate animals: first, in the P0 generation the landing pad strain was crossed to the strain carrying the tag array. F1 male progeny from this cross were then crossed to the recombinase strain. The fraction of mated recombinase-expressing animals that result in a tag varied depending on the array. For C-terminal insertions (UNC-32), about 44% of the mated F1s gave rise to tagged genes for both the GFP and mKate2 arrays; N-terminal insertions (SNB-1) varied between 8 to 24% of mated F1s (Table 3). The tags generated using crosses were viable and expressed the expected fluorescence pattern (Figure 5B).

## Discussion

To understand protein function *in vivo*, the endogenous locus must be tagged to determine when the protein is expressed, where it is localized, and with what other proteins it colocalizes. Here, we present a method using recombinases to insert single-copy transgenes and protein tags into the *C. elegans* genome. A survey of eight recombinases identified six enzymes that act efficiently in the nematode germline, with five being useful in generating insertions.

Combinations of these recombinases can be used for different aspects of genome engineering: A tyrosine recombinase, either Dre or B3, removes plasmid backbone sequences. PhiC31 integrates the construct irreversibly into the genome. Cre recombinase removes the selection marker, and FLP can be used for cell-specific expression. PhIT is particularly useful for modular gene tagging since a palette of fluorescent proteins are available for SapTrap assembly (Figure S2). Moreover, these can be introduced by genetic crosses without the need for difficult microinjections.

Tagged proteins can be generated as multi-copy transgenes, single-copy transgenes, or endogenous tags. PhiC31-mediated integrations can now be used for each of these genome engineering methods: PhIAT (Rich et al., 2025), Single-copy inserts by RMI, RMCE, and PhIT (Beck and Nonet, 2025; Yang et al., 2022), and PhiT tags. The advantage of transgenes, instead of endogenous tags, is that the fluorescence signal can be elevated by increasing copy number or driving expression under strong promoter. The advantage of endogenous tags is that the native context of enhancers and promoters is intact allowing the fluorescence signal to accurately reflect gene regulation. Signal strength has not been a major limitation for endogenous tags for genes tested here, but is expected to be a problem for proteins that are only transiently expressed during development.

The most important features of PhIT are fidelity, efficiency, and modularity. Fidelity relies on the use of recombinases instead of double-strand break repair to insert the tag. Efficiency requires matching the activity of the tyrosine recombinase with the integrase. Modularity allows different tagging cassettes to be inserted into a locus.

### Fidelity

Currently, tagging endogenous genes is performed by CRISPR. The Cas9 endonuclease introduces a double-strand break which is repaired by providing a template that extends across the break. The repair template is copied into the locus by synthesis-dependent strand annealing (SDSA), which is error-prone and non-processive (Frøkjaer-Jensen et al., 2008; Paix et al., 2017; Strathern et al., 1995). Thus insertion of large tags is particularly challenging.

By contrast, integrases, such as PhiC1 (Venken and Bellen, 2012; Yang et al., 2022), can insert large DNA sequences faithfully and without error. Although the enzyme introduces a double-strand break, the DNA is introduced via a regulated and precise crossover of DNA strands. The fidelity of the PhiC31 insertions appears to be quite reliable. Of the several dozen insertions analyzed by PCR, none exhibited damage to the recombinase target sequence. A more serious concern is that long-term exposure of the *attB* site to PhiC31 can damage the target site, which we have observed at an N-terminal *attB* tag of *unc-32* (Figure S8). For this reason, *attB* strains should not be maintained for long periods in the presence of PhiC31 expression.

On the other hand, we observed insertions with corrupted cargo. For example, in one case insertion of the P*npr-2::mKate2* transgene resulted in expression of mKate2 in the intestine and the insertion could not be analyzed by PCR. This error probably occurred during assembly of injected DNA into an extrachromosomal array.

The structure of assembled arrays can also lead to loss of selectable markers. Crossing arrays into the recombinase expressing strain frequently leads to loss of the HisCl channel but the orange array marker remains on the array (Table 3). This specific loss probably arises because array formation is not random (El Mouridi et al., 2022); HisCl plasmids can concatenate by homologous recombination, and this domain is excised by B3 recombinase. For these reasons, we recommend that potential insertions be identified by loss of *all* array markers and confirmed by PCR.

Unlike CRISPR, PhIT tagging of a gene is not scarless. The recombined *attB* site creates *attL* and *attR* sites flanking the fluorescent proteins. The insertion of these flanking may be beneficial. Introduction of linkers between a protein and the fluorescent tag is considered best practice for gene tagging. The PhiC31 sites introduce 14 aa linkers; the tyrosine recombinase will introduce an 11 aa linker. They are in large part determined by the sequence requirements of the recombinase. However, careful consideration was used in choosing the reading frame to avoid potentially troublesome amino acids, and in some cases the overlap sequence of the recombinase was changed to a non-canonical sequence for an optimal linker (Figures S4 and S5). These linkers did not produce notable phenotypes in the genes tagged in this study, and reproduced the expression patterns generated from CRISPR modification of the same genes (Figure S3 (Schwartz et al., 2021)).

PhiT uses CRISPR to insert an *attB* landing pad. Some of the drawbacks associated with CRISPR, such as error-prone repair for long inserts, are minimized: the inserted sequence is only 39 bp, the repair template is an inexpensive single-stranded DNA oligo, and insertion of the *attB* target is highly efficient – insertions are typically observed in most of the injected animals (Ghanta and Mello, 2020; Paix et al., 2017). Nevertheless, about 50% of the *attB*-sized insertions introduced by CRISPR contain errors and must be confirmed by sequencing. Thus, the construction of the *attB* landing pads is currently a limitation for using PhIT for genome-wide gene tagging.

### Efficiency

Recombination efficiencies among the different tyrosine recombinases varied greatly. Our assays were empirical, but we can speculate on causes for differences in efficiency. First, some bacterial recombinases are toxic in eukaryotes; for example, Cre is toxic in *Drosophila* (Nern et al., 2011). Similarly, we found that transgenes for Dre could not be integrated in the *C. elegans* genome. However, Dre can be used for PhIT integrations by injection, in which Dre expression is only transient.

Second, differences in recombinase efficiency could arise from temperature requirements.

Some recombinases are temperature-sensitive, for example, FLP is inefficient in mammals, likely due to low activity at 37°C (Buchholz et al., 1996). In contrast to mammals, FLP is highly efficient in *C. elegans*, possibly because worms are propagated at lower temperatures (15-25°C) (Buchholz et al., 1996).

Third, for PhIT the efficiency of the recombinase needs to match the efficiency of the integrase. In the absence of the tyrosine recombinase, the entire array is integrated into the *attB* site (Rich et al. BioRxiv 2025). If the tyrosine recombinase is less efficient than the integrase the array will be inserted at the tagging site. In one case, we believe a large fragment of an array inserted into the *attB* site for *unc-32*, making the tag lethal if homozygous. Crossing the tyrosine recombinase back into this strain removed this extra sequence, making homozygosity possible. In other cases the recombinase may be too efficient; integration will fail if the array is fragmented before PhiC31 is expressed, as appears to be the case for KD recombinase.

### Modularity

Strains carrying the *attB* landing pad can be used to insert different tags or fluorophores in a modular fashion. Modularity is most important for multicolor imaging, in which colors must be carefully selected to distinguish protein signals. Moreover, the *attB* strains will serve as a resource for the insertion of new tags as they are developed in the future.

One module we include here is cell-specific fluorescent tags. Broad expression creates problems for identifying where a protein is located in a particular cell, since the entire animal will be fluorescent. In these cases, the FLP-on cassette limits protein tagging to a single cell in the absence of an overwhelming cloud of fluorescence coming from other cells. A number of cell-specific drivers of FLP recombinase are available (Fragoso-Luna et al., 2023). A potential concern is that FLP is also used for insertion of single-copy transgenes by the highly efficient method of cassette exchange (Beck and Nonet, 2025; Nonet, 2020). It should be noted that transgenes introduced using RMCE do not preclude use of FLP-on constructs in other tags.

RMCE leaves a single FRT3 site which does not interact with the FRT1 sites used for FLP-on (∼0.5% cross-reaction by FRT3 compared to FRT1)(Schlake and Bode, 1994)

### Genetic Crosses

Importantly, PhIT makes genome engineering broadly accessible. Not all labs have access to the specialized equipment or skills required for microinjection. The strains carrying a specific color on an array, can simply be crossed to an *attB*-tagged gene to generate a tag. The ability to tag proteins in *C. elegans* by crosses will democratize genome engineering of the nematode.

Specifically, it will allow non-specialty labs to test their molecular models in the worm, and encourage schools to conduct genome engineering experiments in the classroom. In the long run PhIT advances the goal of genome-wide gene tagging, so far only achieved for the yeast genome (Huh et al., 2003). The most restrictive limitation now is the number of genes with an *attB* tagging site.

## Methods

### Worm strains and growth conditions

All *Caenorhabditis elegans* strains were derived from N2(Bristol) maintained at room temperature (22°C) on NGM plates seeded with OP50 *Escherichia coli* unless otherwise specified. Strains are listed in Table S2, with some from previous publications (Fragoso-Luna et al., 2023; Frøkjaer-Jensen et al., 2008; Frøkjær-Jensen et al., 2014; Nonet, 2024). Strain construction is described in Extended Data Table ED1.

### Array-based excision assays

Rol reporter generation: The dominant Rol marker (*sqt–1(e1350)*) and assembled by SapTrap using overhanging annealed oligos to generate flanking RT sites (Tables S2 and S3(Schwartz and Jorgensen, 2016)). Each of the Rol excision reporters (RT-Rol-RT) was inserted as a single-copy into chromosome I using either FLP-mediated cassette exchange (Nonet, 2020) or universal MosSCI (Frøkjær-Jensen et al., 2014).

RMCE RT-Rol-RT insertions: Most Rol marker insertion strains were inserted into the *jsSi1570* RMCE site using a custom plasmid (pMR553) (Nonet, 2020). Each RT-Rol-RT plasmid (50ng/µl) was injected into NM5322 (gift from Mike Nonet). After 3 days, Rol F1 animals were picked, and the resulting Rol F2 animals singled and confirmed to be homozygous rollers.

MosSCI RT-Rol-RT insertion: Because the FLP Rol reporter cannot use FLP-mediated RMCE for insertion, universal Mos-mediated single copy insertion was used (Frøkjær-Jensen et al., 2014). The EG8078 (*oxTi185[universal-Mos] I; unc-119(ed3) III)* strain was injected with a mix of 10ng/µL pCFJ1532 Mosase, 10ng/µL pMR564 (FRT::Rol::FRT Cb-*unv-119+)*, 10ng/µL pCFJ104 P*myo-3::mChr*, 10ng/µL pGH8 P*rab-3::mChr*, 2.5ng/µL pCFJ90 P*myo-2::mChr*, 10ng/µL pMA122 *peel-1* negative selection, and 47.5ng/µL Invitrogen 1 Kb Plus DNA Ladder (#10787018) (Extended Data Table 4). After injection, worms were propagated at 25°C for 1 week, and then screened for non-Unc-119 animals that were missing co-injection markers after a 2 hour 20 minute and 34°C heat-shock. These animals were further screened nonUnc homozygous Rol (*FRT::sqt-1::FRT*) animals lacking co-injection markers, indicating presence of an insertion and not a persistent extrachromosomal array.

Excision assay. Each Rol reporter strain was injected with reagents for pharyngeal red fluorescent (P*myo-2*::mCherry) array marker (2.5ng/µL) and germline expressed (P*mex-5*) recombinases (12.5ng/µL). We also injected DNA ladder as stuffer to promote array formation (85ng/µL). F1 Rol animals bearing the red fluorescent extra-chromosomal markers were singled and maintained at 25°C. These worms were transferred to new plates every day for three days. After 3 days of growth F2s were assayed for the presence of Rol phenotype. Only F1s that had any number of non-Rol phenotypes were recorded, since it was unclear if F1s without non-Rol F2s received the recombinase. The frequency of excision was calculated as follows:

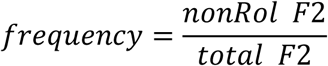

### Recombinase cross excision assays

Recombinase expression constructs were integrated into the genome by FLP-based cassette exchange and MosSCI (Frøkjær-Jensen et al., 2014; Nonet, 2020) on chromosome II, except for *jsSi1623* which was a gift from Mike Nonet and is found on chromosome IV.

Recombinase insertion by RMCE. Most chromosome II recombinase insertion strains were generated by recombinase mediated cassette exchange (Nonet, 2020). 50ng/µl of each plasmid were injected into NM5304 (gift from Mike Nonet). After 3 days, Rol F1 animals were picked, and allowed to produce offspring. Rol F2 animals were singled and allowed to reproduce. When the progeny of these animals appeared homozygous for insertions with 100% Rol progeny and lacking the GFP landing pad maker, L2 animals were heat-shocked at 34°C for 2 hours and 20 minutes. After heat shock, these animals were allowed to reproduce, and wild-type offspring were selected.

MosSCI insertion of Cre. EG6699 (*ttTi5605 I; unc-119(ed3) III*) was injected with a mix of 10ng/µL pCFJ1532, 10ng/µL pCFJ104, 10ng/µL pGH8, 10ng/µL pMA122, 2.5ng/µL pCFJ90, 47.5ng/µL Invitrogen 1 Kb Plus DNA Ladder (#10787018), and 10ng/µL pMR547 (Extended Data Table 4). After injection, worms were allowed to reproduce at 25°C for 1 week. After 1 week, plates were screened for non-uncoordinated animals that were missing co-injection markers. These animals were screened for insertions by nonUnc animals with green germlines lacking co-injection markers.

Cross-based excision assay. To determine the excision efficiency of single-copy insertion recombinases, males homozygous for these transgenes and heterozygous for a chromosome I balancer (*tmC27* (Dejima et al., 2018)) were crossed with the Rol reporter strain. These strains were allowed to mate at 20°C for 2 days before transferring P0 hermaphrodites to new plates. The balancer carries a mutation in *unc-75* and a pharyngeal Venus expression construct, and balances the excision marker, which would otherwise be invisible. Rol F1s with fluorescent green pharynxes were singled 3 days later and were transferred every day to new plates for 3 days. These animals were heterozygous for the recombinase Rol marker balanced by *tmC27*. Three days after transfer, F2s were scored, where Unc worms were homozygous for the balancer, non-green (bal -) worms were homozygous for the marker, and green non-Unc worms were het for the excision marker. The latter group was used in our calculations for excision frequency, as follows:

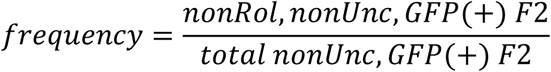

### Generating *attB* landing pads

CRISPR design and mix preparation. RNP CRISPR injection mixes contain tracrRNA, crRNA, and single-stranded oligodeoxynucleotide (ssODN, ultramer) oligos from IDT (Dokshin et al., 2018). crRNA and tracrRNA were diluted with nuclease free duplex buffer (IDT) to a concentration of 100µM and 18µM respectively. crRNA guides were chosen based on proximity to the desired insertion site, where the closest cut-site possible was selected. The ssODNs were diluted to a concentration of 100µM with Invitrogen UltraPure Water (#10977015). Repair templates with 50 bp homology arms were designed to edit the recognition site or PAM sequence, depending on which was in the same direction from the cut as the desired insertion. To anneal the RNA oligos, 2.5µl of crRNA was added to 5µL of tracrRNA, they were then heated to 95°C for 5 minutes, before air-cooling for 5 minutes while keeping the tube inverted to prevent rapid cooling by conduction. Next, 1.3µL of Cas9-nls (Berkeley, MacroLab) was added, and allowed to incubate at room temperature for 5 minutes. Finally, 1µL of ssODN was added along with 500ng of pSEM233 (final concentration of 25ng/µL) and ultra-pure water to a final volume of 20µL.

CRISPR injection. Before injection, the CRISPR mix was spun at 10,000 rcf for 10 minutes to ensure debris is pelleted. Young-adult Unc worms (EG9814, *unc-119(ox819)*) were injected and maintained at room temperature for 3 days, after which, F1 worms with the red co-injection marker (pSEM233) were singled and allowed to produce offspring before being lysed and genotyped as described below.

Genotyping PCR. After injecting animals with a CRISPR mix for generating an *attB* insertion, we lysed and PCRed up to 16 F1 animals that had a fluorescent co-injection marker, after laying eggs. Genotyping PCR was done as described in Wormbook (Fay, 2013). Single-worms were lysed in 10 µL of a fresh mixture of worm lysis buffer using proteinase K obtained from New England Biolabs (NEB #P8107S). Each PCR reaction contained 7.24 µL water, 1 µL Taq buffer (NEB #B9014S) 0.2 µL 10mM dNTPs (Thermo Fisher, #R0193), 0.2 µL 10mM forward primer, 0.2 µL 10mM reverse primer, 0.1 µL 50mM MgCl_2_(NEB #B0510A), and 0.06µL Taq DNA polymerase (NEB #M0273). All primers were obtained from Integrated DNA Technologies. The PCR cycling conditions were: 95°C initial denaturation for 1 minute, followed by 30 cycles of 95°C for 25 seconds, 20 seconds of 56°C for annealing, and extension at 68°C for 1 minute/kb. Using these amplicons, we used gel electrophoresis with a 2% agarose (GoldBio and TAE (Tris-acetate-EDTA) buffer) to identify gel shifts that indicated the presence of an insertion. When homozygous for the insertion, Sanger sequencing (Azenta) was used to determine the insertion sequence; precise sites are indicated (Table S1). We found that on average about 50% of appropriately sized insertions had the correct *attB* sequence.

### PhIT integration by injection

To generate gene tags and transgenes by injection, young adult Unc worms (*unc–119(ox819) III*) with genomic *attB* tag sequences were injected. The *ox819* allele was generated in a wild-type background to ensure an unmutagenized genetic background (Schwartz and Jorgensen, 2016). The injection mix contained either plasmid or linear DNA in the following concentrations: 10ng/µL pWD540 (P*eft-3::LSSmOrange* array marker), 50ng/µL tagging plasmid, 20ng/µL pAJH98 (*Pmex-5::PhiC31*), 10ng/µL pAJH16 (Dre) or pAJH15(B3), and 10ng/µL Invitrogen™ 1 Kb Plus DNA Ladder (#10787018). After injection, worms were transferred to an OP50-seeded NGM plate for recovery. All injected P0s were maintained together on one recovery plate at 25°C. Three days after injection, wild-type F1s were singled onto OP50 seeded NGM plates. 5 days after singling, plates were examined for wild-type F2/F3 lacking the orange array marker. To confirm tag insertion, we singled wild-type F2/F3 lacking array markers, confirmed mendelian inheritance of wild-type rescue (100% or 75% wild-type), identified the correct fluorescence pattern, and performed PCR as described for *attB* insertion to confirm insertion. The primers used are listed in Extended Data Table 2. Tagging efficiency was determined by the percent of F1 plates that had any number of tagged F2 progeny.

### PhIT tagging by crossing

Array strains carrying the desired tag were generated by injecting plasmid DNA in the following concentrations: 10ng/µL pWD540 (P*eft-3::LSSmOrange* array marker), 50ng/µL tagging plasmid, 20ng/µL pMNK56 (*him-8* piRNA), 10ng/µL pSEM238 (HisCl1, histamine chloride channel), and 10ng/µL Invitrogen™ 1 Kb Plus DNA Ladder (#10787018). *him-8* piRNAi was included to produce Him (high-incidence of males) animals that would facilitate easy mating of strains (Priyadarshini et al., 2022).Tag array-bearing males were crossed to *unc–119(-)* strains carrying the *attB* landing pads. Wild-type F1 animals from this cross carry both the tag in the array and the target *attB* landing pad but lack the recombinase enzymes (Figure 5A). F1 males were then crossed to *unc–119(-)* PhiC31 and B3 expressing animals (EG10391). The mated recombinase-expressing animals were singled, and allowed to reproduce for 10 days at 25°C. Because the *attB*-tagged locus, the fluorescent tag, and the recombinases are all present, the array is fragmented and the tags are inserted into the *attB* site. After 10 days, plates were treated with histamine, which paralyzes animals still carrying arrays expressing the histamine-gated chloride channel. Insertions were detected by presence of wild-type locomotion, absence of the orange array marker, presence of the expected fluorescence, and confirmed by PCR.

Tagging efficiency of crosses was determined by: the fraction of mated recombinase-expressing F1 animals that result in a tag (Figure 5A, Table 3) (genotype: *attB or + /+; PhiC31-B3/ +; unc-119(-); oxEx*[fluorescent tag]). Note that histamine-resistance was not a reliable read-out for loss of the array; histamine-resistant F4 animals can sometimes arise because B3 recombines the array and sheds the region containing the histamine-sensitive Cl channel.

### Plasmid Assembly

All plasmids generated or used in this study are listed in Extended Data Table 3. ApE-formatted sequence files for all plasmids generated in this study are available as supplemental data. Some plasmids were gifts from Mike Nonet, Christian Frøkjaer-Jensen, and Joachim Messing (El Mouridi et al., 2021, 2020; Nonet, 2020; Priyadarshini et al., 2022; Vieira and Messing, 1991).

Golden Gate donors were generated by restriction digest and blunt-end ligation of PCR products or double-stranded synthetic DNA (see Extended Data Table 2). PCR and restriction fragments were column purified or gel extracted (QIAGEN) before ligation cloning. Restriction digest cloning reactions had the following: 1µL 50ng/<u>µ</u>L destination vector, 50ng/µL or less of PCR amplicon, 1µL rCutSmart Buffer (NEB, B6004), 1µL restriction enzyme(s) (NEB, see Table), 5µL T4 Ligase (NEB, M0202), 1µL 10mM (NEB, P0756), and water to a final volume of 10µL. After mixing, the reactions were placed in a 25°C incubator overnight. The following day, these reactions were heated at 65°C for 30 minutes, then re-treated by the cloning restriction enzyme before transforming into DH5α competent *E. coli* cells (NEB 5-alpha).

Amplicons used in cloning were generated with 2X Q5® Hot-Start Master Mix from NEB (#M0494). Manufacturer’s recommendations for extension time, amount of template DNA, and annealing temperature were followed. See Extended Data Table 2 for a list of primers used to generate amplicons used in cloning.

B3, Dre, KD, and Vika coding sequences were codon optimized using the webtool developed by Redemmen and colleagues (Redemann et al., 2011). Three synthetic introns were included: the *rpl-18* intron 2, GG3 for use in cloning PATC introns (Aljohani et al., 2020), and an intron generated by the webtool, respectively. SapI sites were incorporated at the extreme 5’ and 3’, while additional SapI and BsaI restriction sites were removed by synonymous codon changes. These sequences were purchased from Twist Bioscience™ as Kanamycin resistant plasmids (pAJH1-4). To ensure germline expression, the PATC intron 3 from *smu-2* (Aljohani et al., 2020) was added to each coding region at the second intron position using pCFJ1149 and BsaI Golden Gate assembly. Using SapTrap reactions, the resulting germline optimized recombinase coding sequence plasmids were assembled with an RMCE destination vector (pNM5340, gift from Mike Nonet), the *pie-1* promoter (pAJH8) and *SL2::mNeongreen::glh-2* 3’UTR (pAJH10) in a ‘SapTrap’ reaction.

Except for the FLP excision reporter (pMR564), recombinase reporters were generated using SapTrap assembly of annealed overhanging oligos (Extended Data Table 2), pMR553 (RMCE destination vector), and pMR551 (*sqt-1(e-1350)* donor).

Most tag and transgene plasmids for PhIT were generated by SapTrap (Schwartz and Jorgensen, 2016). The standard 2.5µL SapTrap reaction (Schwartz and Jorgensen, 2016) was assembled for SapI Golden Gate. For BsaI golden gate we used a 10 µL mix containing 1 µL BsaI-HF®v2 (NEB®, R3733), 1 µL CutSmart® buffer (NEB® B7204), 1 µL 10 mM ATP (NEB® P0756), 0.5 µL T4 ligase (NEB®, M0202) and 0.05 pmols of plasmids, using water to achieve a 10 µL final volume. For PaqCI cloning of pAJH93, a 20 µL mix of 2 µL T4 ligase buffer (NEB® B0202), 2 µL T4 ligase (NEB®, M0202), 2 µL PaqCI (NEB® R0745), 2 µl PaqCI activator (NEB® S0532), 0.1 pmol of each plasmid, 0.3 pmol of annealed oligos, and water was assembled. After assembling each mix, reactions were placed overnight at 25°C, and heat-killed and re-treated with the respective cloning enzyme the next day before transformation into NEB® 5-alpha cells (C2987). See Extended Data Table 3 for plasmid details and Extended Data Table 2 for oligo sequences.

Phit transgene destination vectors for B3, Dre, and KD injections (pAJH83, pAJH120, and pAJH122 respectively) were generated by combining pAJH42, pMLS253(*lox*-flanked *Cbr unc-119* rescue) and overhanging oligos (Extended Data Table 2) to generate a multiple cloning site and appropriate recombinase targets. After cloning these constructs, a new SapI landing pad was generated by restriction digest cloning with annealed overhanging oligos (Extended Data Table 2). With SapI sites incorporated, we combined these plasmids with promoter, 3’UTR, and CDS plasmids in a SapTrap reaction. For pAJH147, which has PhiC31 and B3, we performed sequential rounds of Golden gate assembly, where *Pmex-5::PhiC31::SL2::mNeonGreen::glh-2* was assembled by SapI golden gate into pAJH83 to generate pAJH103, after which we cut and added SapI sites by restriction digest with NotI (NEB® R0189) and annealed overhanging oligos (pAJH110). Finally Golden Gate assembly was used to assemble *Pmex-5::B3::let-858* into the plasmid, generating pAJH147.

Phit tags were generated by SapI Golden Gate assembly into pAJH42. pAJH42 was generated by cutting Litmus28 (Evans et al., 1995) with NcoI (NEB® #R103) and ligating annealed overhanging oligos into the cut site. Care was taken to generate oligos which produce tags that would have an in frame a recombinase target linker sequence (N-terminal and internal tags) or a premature stop before the recombinase target scar (C-terminal tags).

### Imaging

Images stacks were acquired using a Zeiss LSM 880 AxioObserver confocal microscope and 1.5 thickness coverslips. Young adult worms were imaged 21-24 hours after picking L4s, and were maintained at 20°C. For tagged SNB-1 and UNC-32 strains, worms were mounted for imaging by using a 2% agarose pad on a glass slide, paralyzed with a combination of 2.5µL of 180mM muscimol (Sigma # M1523) and 2.5µL of polystyrene beads (Sigma #84135). Images are then processed by FIJI (Schindelin et al., 2012) where max intensity Z-projections were generated and overlayed with a single slice from the mock-DIC channel.

LAAT-1::FLPon::Skylan-S (Figure 3): Images were taken with a 40x Plan-Apochromat 1.3 Oil DIC UV-IR M27 objective. Z-stacks were acquired through the whole worm using a 2 µm step interval. Individual channels were imaged using sequential illumination with 405 nm light to capture autofluorescence (switching on Skylan-S) and 488 nm light to capture fluorescent signal from Skylan-S (which also switches off Skylan-S).

mKate2 transgenes (Figure 2): Images stacks were acquired with a 40x Plan-Apochromat 1.3 Oil DIC UV-IR M27 objective. Z-stacks were acquired through the whole worm using a 3 µm step interval. Individual channels were imaged using sequential illumination with 405 nm light to capture DIC (transmitted light) and 594 nm light to capture fluorescent signal from mKate2.

Note: All mKate2 images except *Pdat-1::mKate2* chr I (upper left panel) were collected using the same number of pixels (4660 x 4660) and same field of view (354.25 µm x 354.25 µm). *Pdat-1::mKate2* chr I was collected using the same number of pixels (4660 x 4660) but a smaller field of view (224.92 µm x 224.92 µm), resulting in a higher pixel density.

### 5-phenyl auxin synthesis

To a solution of 2-(5-bromo-1*H*-indol-3-yl)acetic acid (5g, 19.7 mmol, 1 equiv.) in dry methanol (100 mL), was added to acetyl chloride (7.0 mL, 98.5 mmol, 5 equiv.) dropwise at 0 °C. The solution was warmed to RT over 3h at which time complete conversion was determined by TLC. The solution was then condensed to ∼10 mL and added to a separatory funnel and neutralized with sat’d NaHCO_3_. The solution was extracted with 3 x 15 mL DCM and the combined organic fractions were washed with brine, dried over Na_2_SO_4_ and condensed by rotary evaporation to afford 5.1 g methyl 2-(5-bromo-1*H*-indol-3-yl)acetate as an off-white powder. The material was carried on without further purification.

In a 40mL vial equipped with a stir bar and septum cap was successively added Pd(OAc)_2_ (25 mg, 0.11 mmol, 0.01 equiv); SPhos (91 mg, 0.22 mmol, 0.02 equiv), anhydrous K_3_PO_4_ (4.8 g, 22.4 mmol, 2.0 equiv), PhB(OH)_2_ (2.1 g, 16.8 mmol, 1.5 equiv.), and methyl 2-(5-bromo-1*H*-indol-3-yl)acetate (3.0 g, 11.2 mmol, 1.0 equiv.). The vial was then sealed and briefly shaken to mix the solids. The atmosphere was replaced 3x with vacuum/nitrogen cycles, and dry degassed toluene (25 mL) and the solution was heated to 100 °C for 4h. The solution was filtered hot through a pad of celite and the filter cake was washed with 10 mL EtOAc. The combined filtrate was then condensed and purified by column chromatography (10% → 50% EtOAc/Hexane) to afford 2.6 g of methyl 2-(5-phenyl-1*H*-indol-3-yl)acetate (87.5% yield) as a white powder.

To a solution of methyl 2-(5-phenyl-1*H*-indol-3-yl)acetate (2.6 g, 9.8 mmol, 1 equiv.) in methanol (20 mL) cooled to 0 °C was added KOH (10 mL, 1M in methanol). The solution was stirred until complete by TLC then acidified to a pH ∼3 by addition of Dowex-H^+^. The solution was filtered and condensed by rotary evaporation then dried overnight at RT to afford 2.4 g (98%) of the title compound as a white powder. ^1^H NMR (300MHz, DMSO-*d*_6_) 𝛿: 3.14 (s, 2H), 3.73 (s, 2H), 7.30 (d, 4H), 7.45 (m, 4H), 7.65 (d, 2H), 7.80 (s, 1H), 11.00 (s, 1H), 12.20 (1H). Spectrum matches known compound (Figure S7).

---

**Figure S1:**
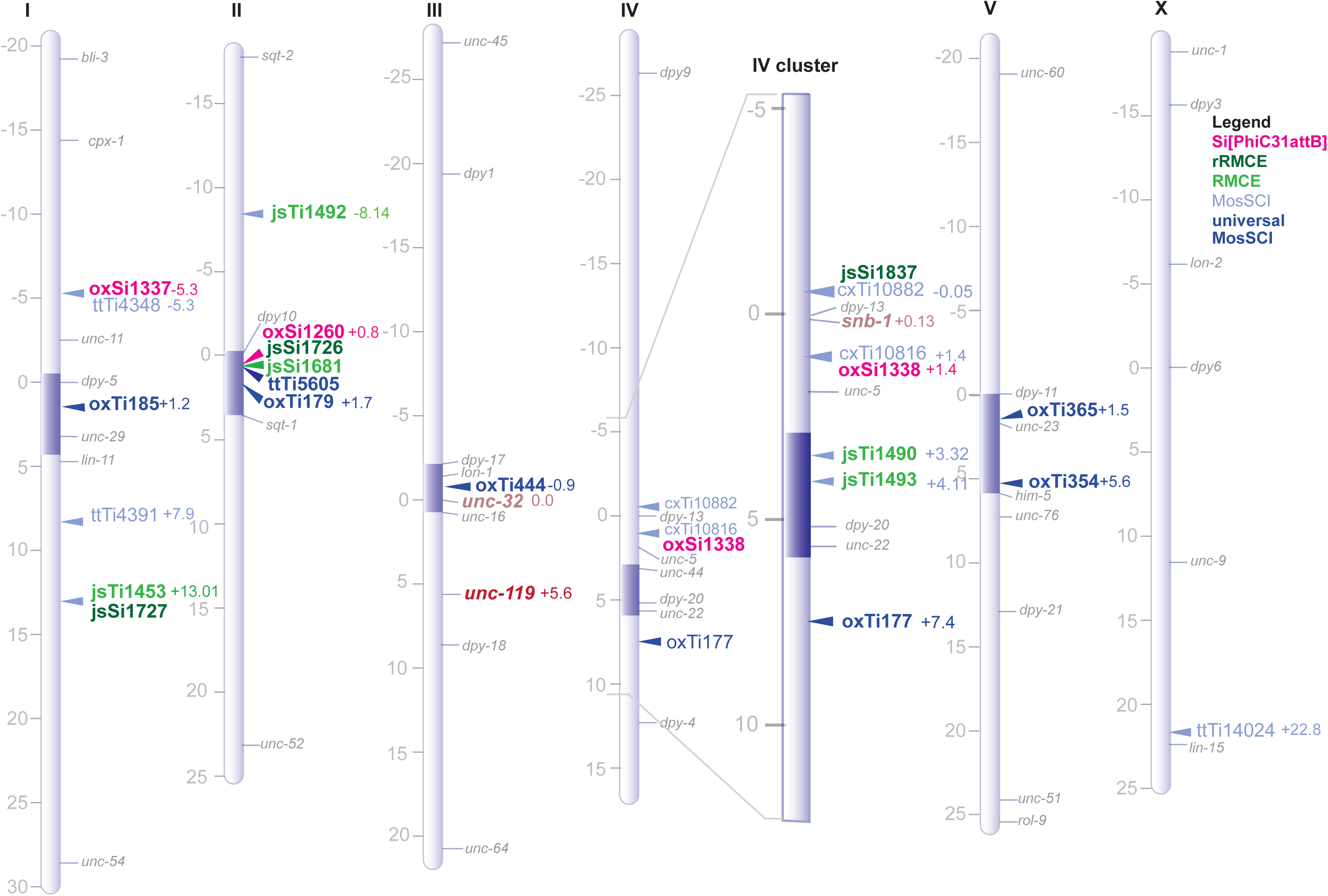
Genetic map of *attB*, MosSCI, and RMCE sites. *attB* landing pads were integrated by CRISPR (pink). Dark and light green indicates locations of rRMCE and RMCE sites, respectively (Nonet, 2023, 2020). Light blue and dark blue represent sites for MosSCI and universal-MosSCI respectively (Frøkjaer-Jensen et al., 2008; Frøkjær-Jensen et al., 2014). Genetic position indicated in centimorgans.

**Figure S2:**
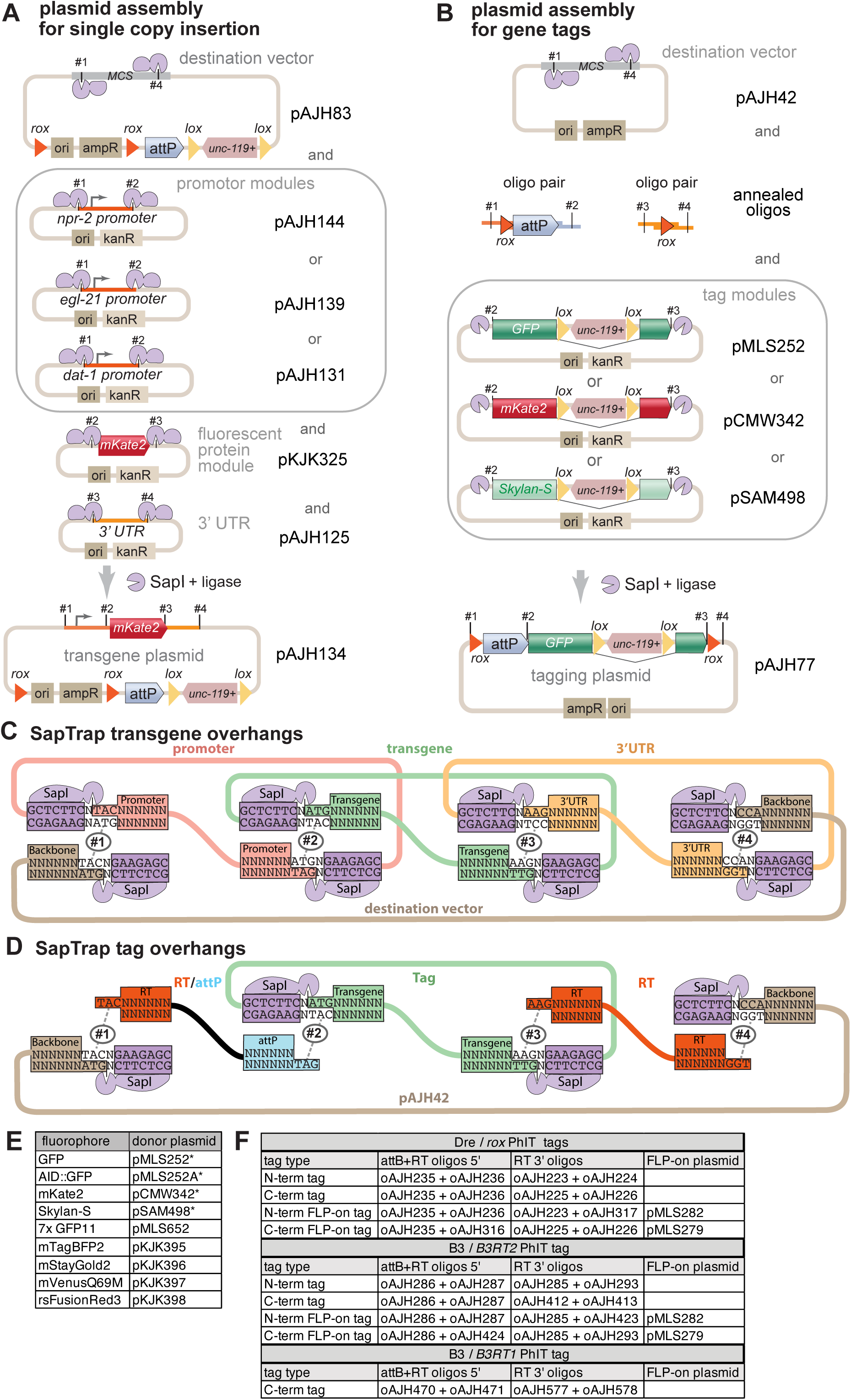
SapTrap assembly. (A) Transgene assembly for single copy insertion. The destination vector contains both the PhiC31 *attP* site for integration and two tyrosine recombinase sites for excision of backbone sequences (for example *rox* sites that are targeted by Dre recombinase, as shown). The destination vector also contains an *unc–119(+)* rescuing transgene as a selection marker; the *unc–119* gene is flanked by *lox* sites so that it can be removed from the transgene by Cre recombinase. Transgenes in this assay were assembled by mixing three donor vectors – a promoter, fluorescent protein, and *let-858* 3’ UTR – with the destination vector and SapI enzyme for one pot DNA assembly. Alternative donor plasmids can be used as interchangeable modules to create constructs with different promoters, fluorescent proteins, and 3’ UTRs. (B) Gene tag assembly. The destination vector pAJH42 has SapI sites for SapTrap cloning. PhIT tags in this paper were assembled by mixing the destination vector pAJH42 with two pairs of annealed overhanging oligos and a fluorescent protein donor vector which has an intronic *unc–119* rescue in a SapTrap assembly mix. These overhanging annealed oligos encode for the RT and *attP*, or just the RT sequence. The fluorescent protein donor plasmids are interchangeable, using the same SapI sites, simplifying tag construction. (C) SapTrap assembly using plasmids. Overhanging bases generated by SapI digest are labeled with numbers (#N) to show complementarity and annealing during assembly. (D) SapTrap assembly using overhanging oligos. Overhanging bases generated by SapI digest or annealing of oligos are labeled with numbers (#N) to show complementarity and annealing during assembly. (E) Fluorescent protein tags available. The fluorescent proteins all contain an *unc-119+* rescuing marker in an intron flanked by lox sites for excision by Cre recombinase. These tags all utilize the #2 and #3 Sap1 sites for SapTrap assembly as shown in the examples above. Asterisks indicate those plasmids used in this manuscript. (F) Oligos for generating recombinase target and attB or just the recombinase target. C–terminal recombinase targets encode a stop codon before the target. For FLP-on constructs, the plasmid used for FLP-on is indicated.

**Figure S3:**
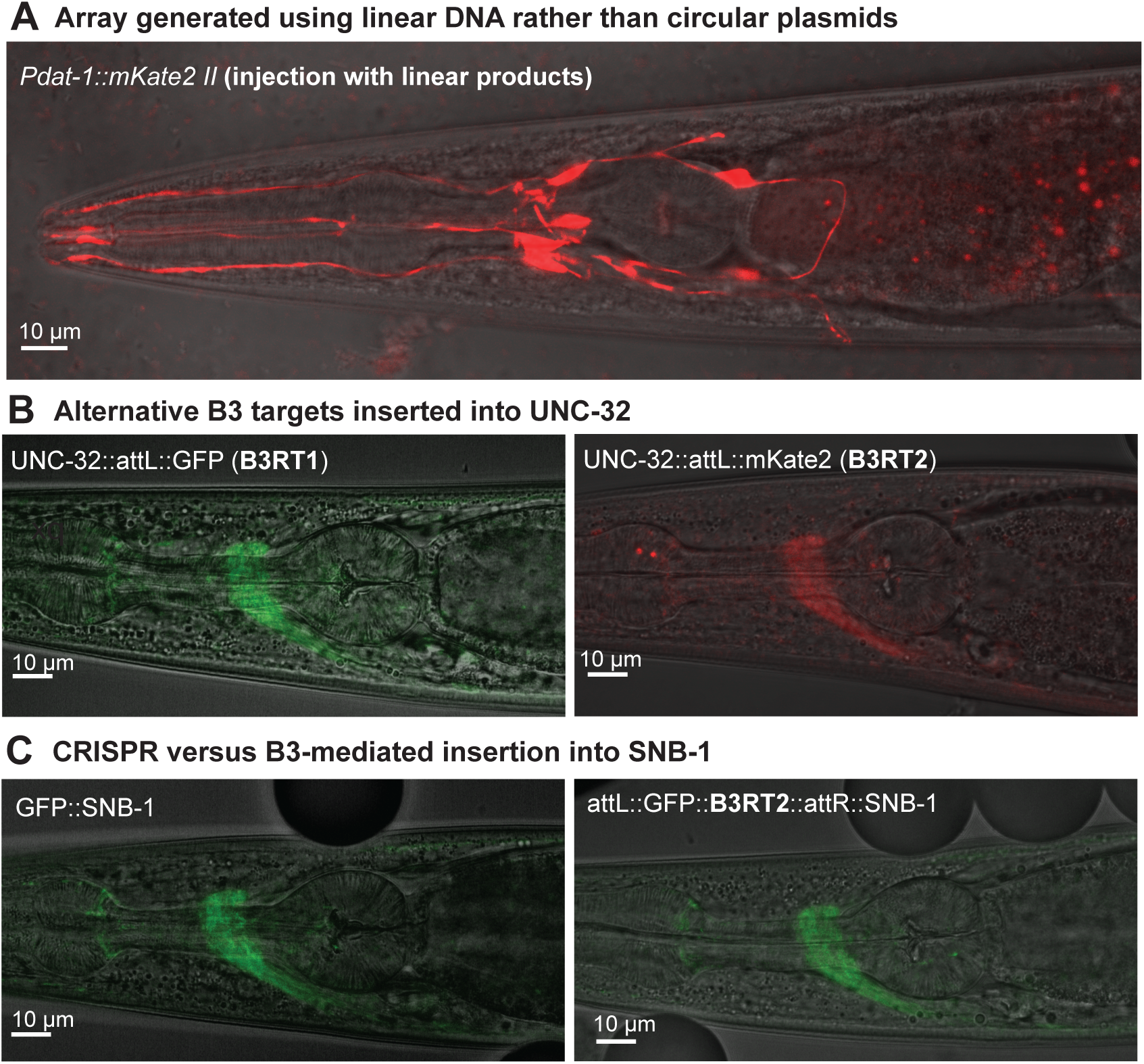
Additional images of strains. (A) Single-copy transgene insertion made by injection of linear DNA instead of circular plasmids. Restriction enzyme digested plasmids were injected at the same concentrations as circular plasmid reagents into the chr II *attB* landing pad strain (EG10204). Linearized DNA and circular DNA generated integrations at the same frequency and same expression pattern as circular DNA for P*dat-1::mKate2*. (B) UNC-32 tagged with GFP or mKate2 tags using *B3RT1* or *B3RT2* sites, respectively. Two examples of B3-mediated tags in UNC–32 made by injection. Expression patterns were indistinguishable from tags bearing *rox*-sites. (C) SNB-1 tagged by CRISPR or tagged by PhIT. The expression pattern of GFP-tagged SNB-1 generated by CRISPR (left), which lack scars left by recombinase target sites (Schwartz et al., 2021), was similar to a GFP-tagged SNB-1 strain generated by PhIT using *B3RT2* sites (right, EG10429)

**Figure S4:**
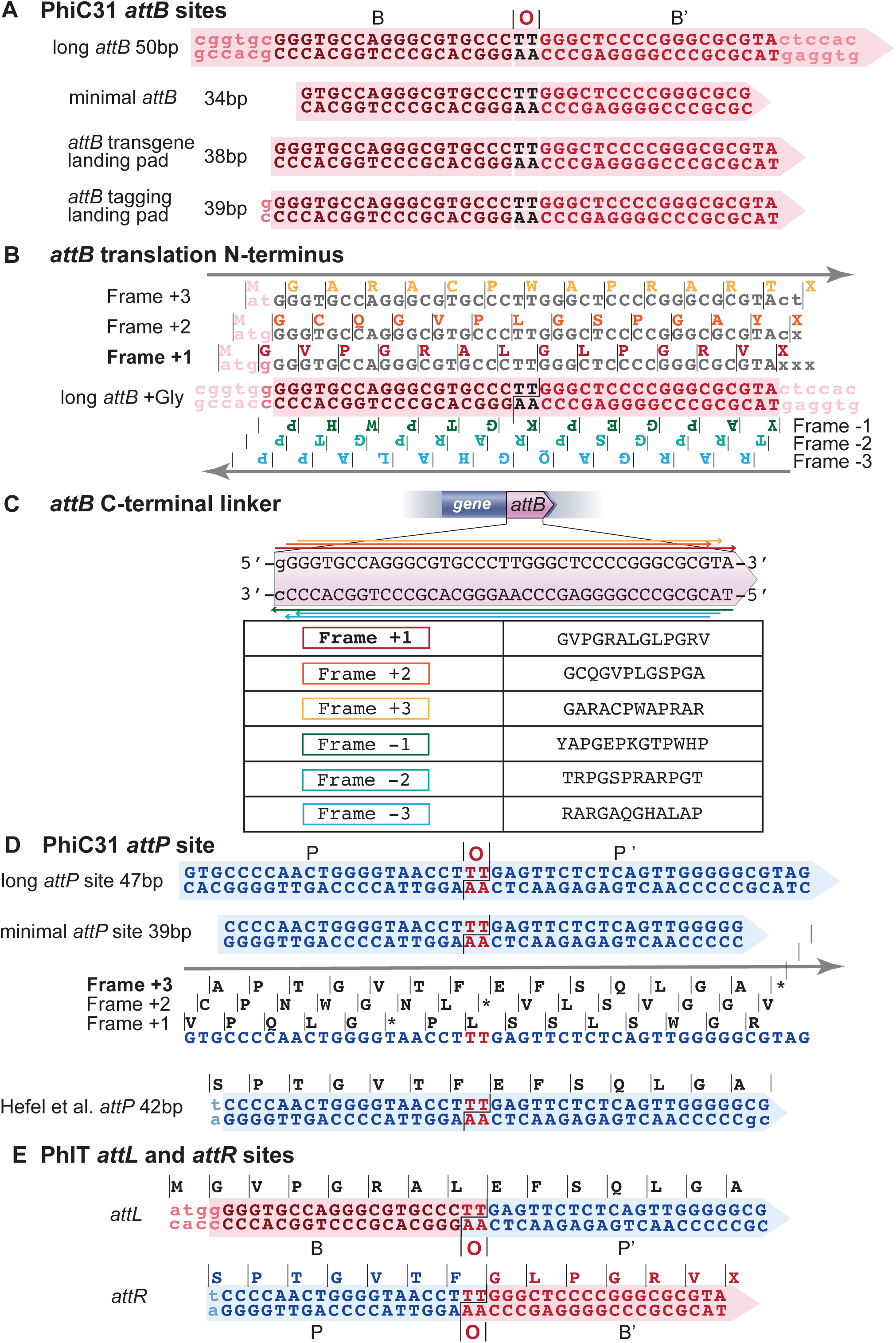
PhiC31 *attB* and *attP* linkers. (A) PhiC31 *attB* sites. The bacterial attachment site(*attB*) for the phage PhiC31 was characterized in the bacteria *Streptomyces lividans* (Groth et al., 2000; Gupta et al., 2007) . These sites are pseudo-inverted repeats (B and B’) centered around an overlap (O) sequence at the cut site. *attB* sites between 34bp to 50bp are functional. (B) All six reading frames were compared to select a functional *attB* site that could act as an N-terminal extension for a tagged protein The forward strand was selected because all three reading frames would be translated as a Met•Gly pair which are highly stable as N-terminal residues (Gonda et al., 1989). (C) Reading frames were compared to select a C-terminal linker between the protein and a fluorescent protein. The +1 frame generates a 13aa linker. (D) The phage attachment site (*attP*) was translated in the three forward reading frames to match the *attB* site. The +3 frame was selected because it does not introduce a stop codon and was optimized to interface with a tyrosine recombinase at its N-terminus. (E) After integration the inserted tag will generate a chimeric site on the left (*attL* BOP’ site) and a chimeric site on the right (*attR* POB’), which form the new linker regions in the tagged protein.

**Figure S5:**
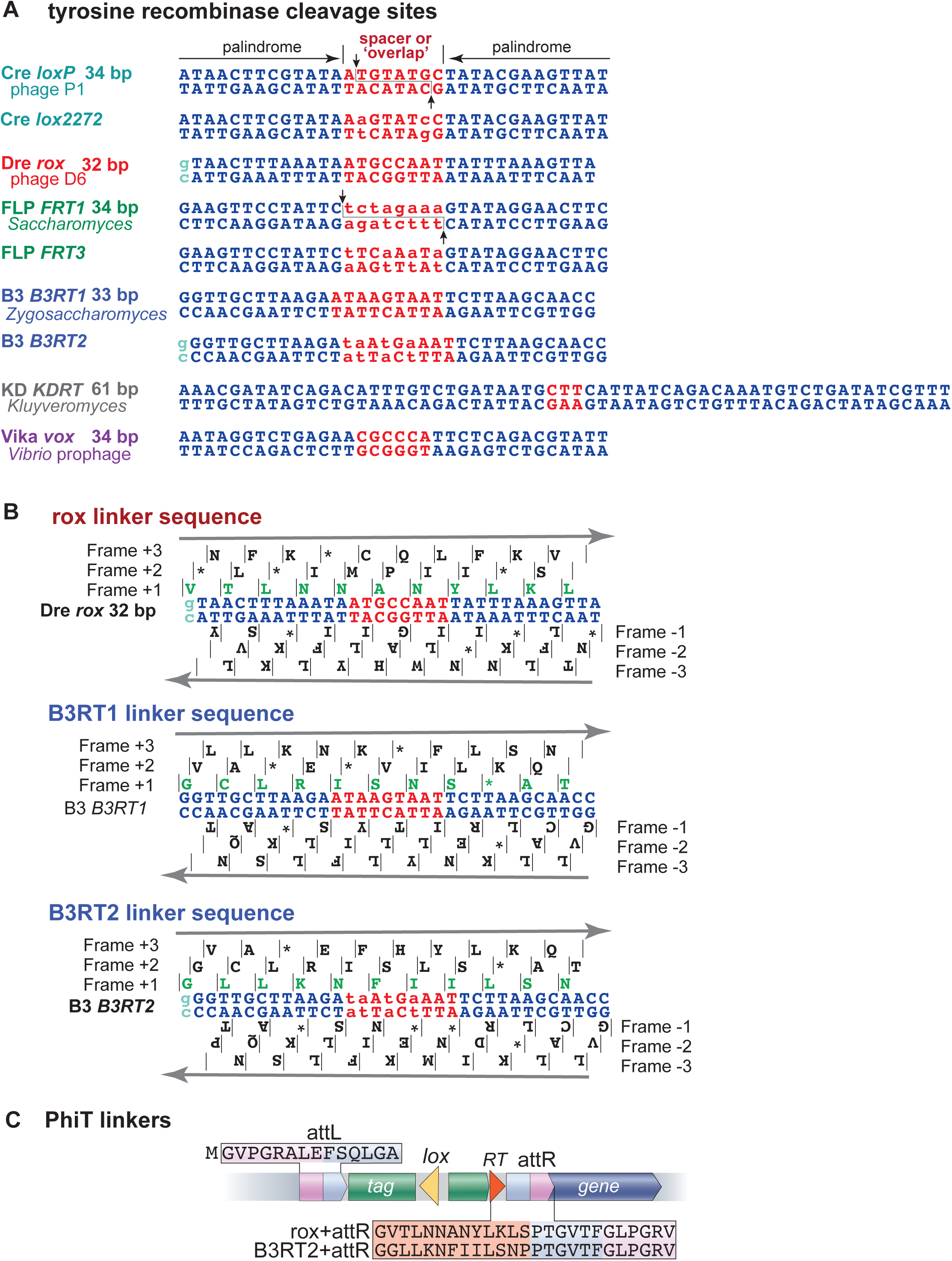
Tyrosine recombinase targets and linkers. (A) Recombinase target sequences. Tyrosine recombinase targets contain inverted repeated of palindromic sequences that bind the recombinase (blue). These sequences flank a spacer, or ‘overlap’, region (red) which has few interactions with the recombinase (Meinke et al., 2016) and define directionality and specificity of recombination. Nucleotides in the overlap that differ from the canonical sequence are shown as lowercase letters. In addition, basepairs were added to the selected open reading frame (bold) to avoid stop codons and optimize linker sequences (teal). (B) Possible translations of target sequences. All recombinase targets used to tag genes are shown with each possible reading frame. The selected sequence and reading frame are all +1 (green). (C) Translation outcomes of PhIT linker scars. In genes with an N–terminal tag, the attR linker is preceded by either a *rox* or *B3RT2* recombination target. For C–terminal tags, the tag contains the stop codon, so that the tyrosine recombinase and *attR* sites are not incorporated into the protein.

**Figure S6:**
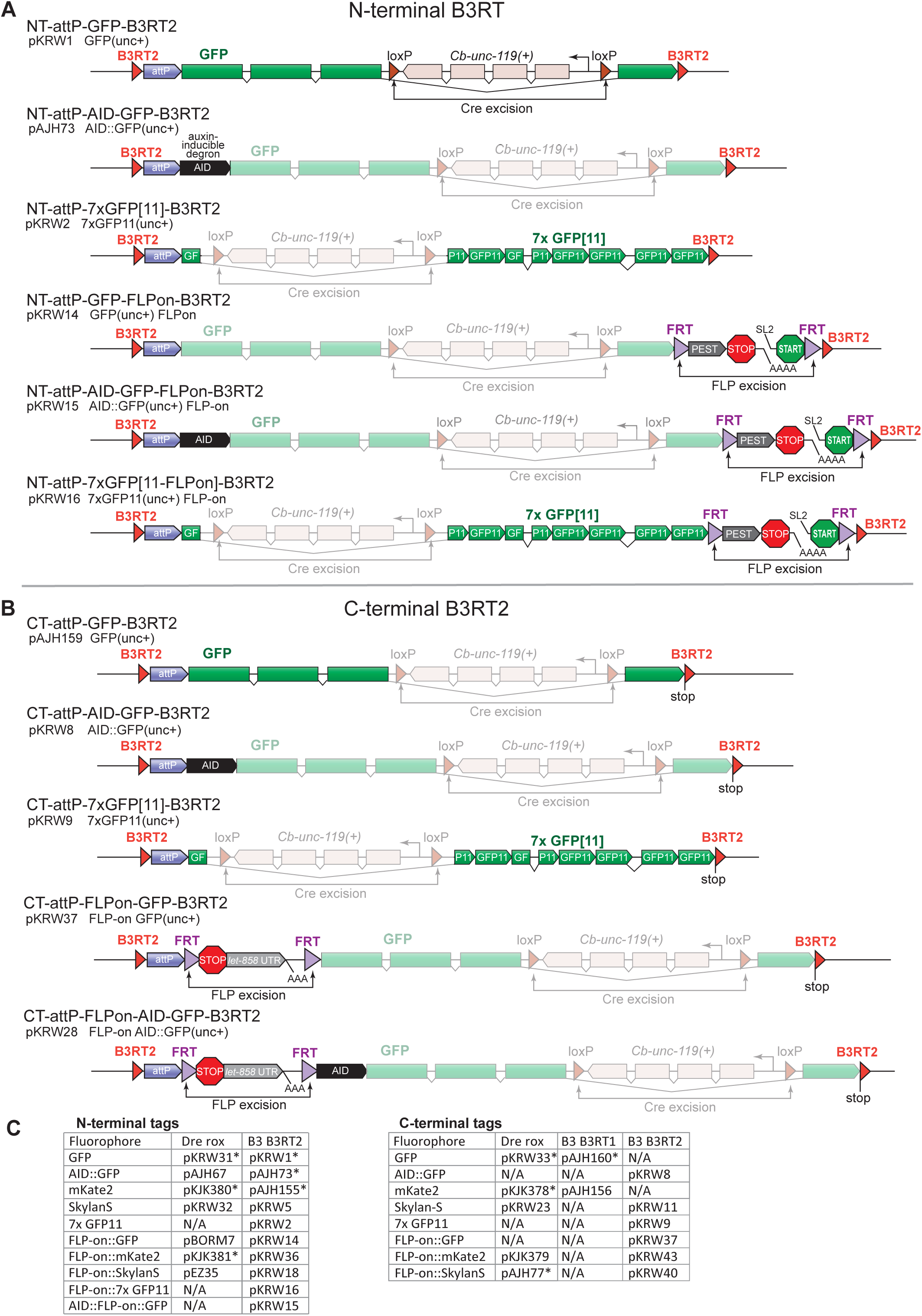
PhIT Modules. (A) N-terminal tags. Examples of N-terminal PhIT tags using B3 recombinase target *B3RT2*. (B) C-terminal tags. Examples of C-terminal PhIT tags using B3 recombinase target *B3RT2*. (C) Tag plasmids list. All tags contain an *unc-119(+)* rescuing transgene in an intron of the fluorophore as illustrated above. Asterisks indicate plasmids used for this publication.

**Figure S7:**
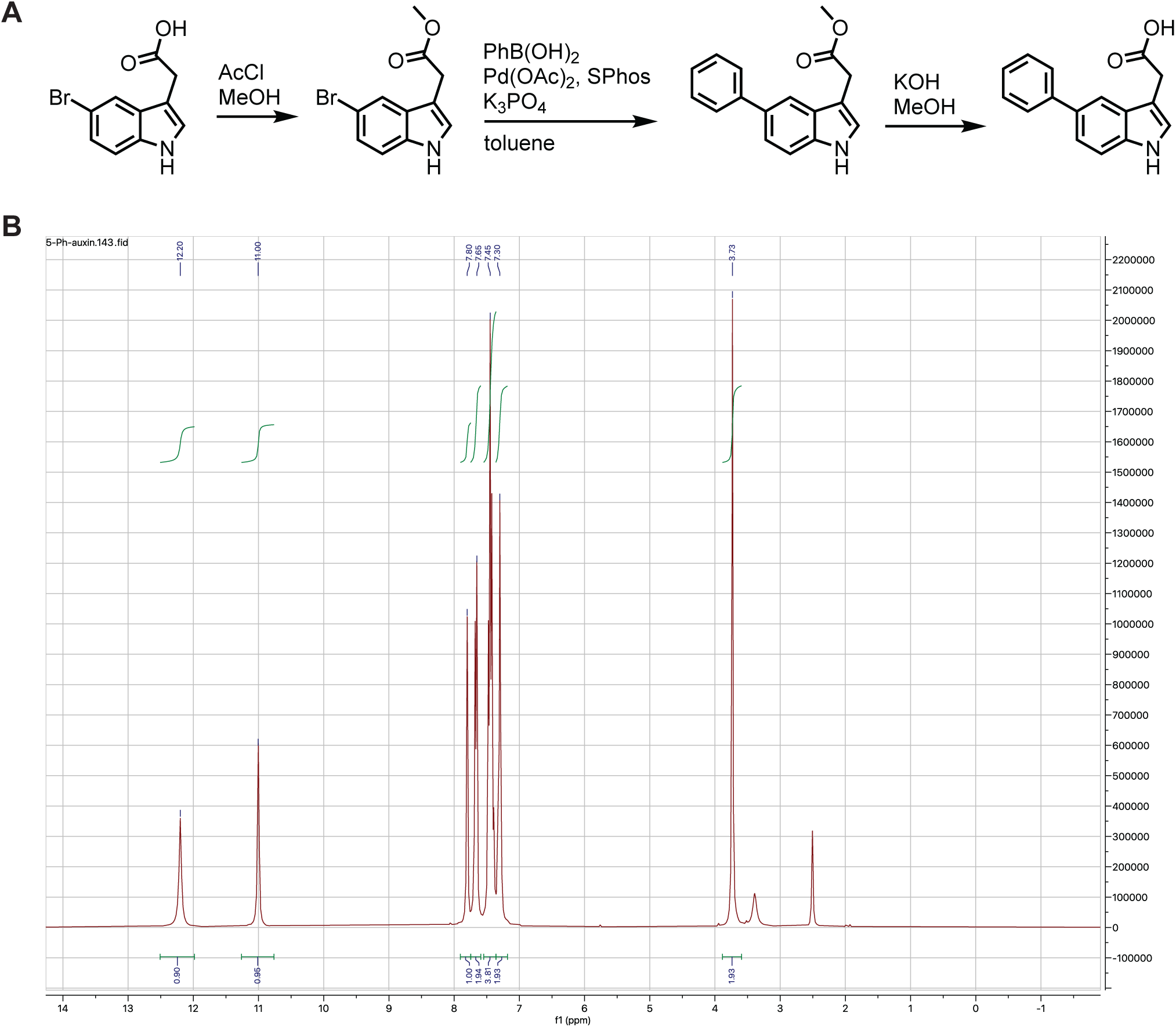
5-phenyl auxin synthesis. (A) Chemical synthesis of 5-phenyl auxin (see Methods). (B) NMR spectrum for yielded product. Spectrum matches known compound. Although the starting reagents differ from a previously published method, final yield and purity are similar (Sural et al., 2024).

**Figure S8:**
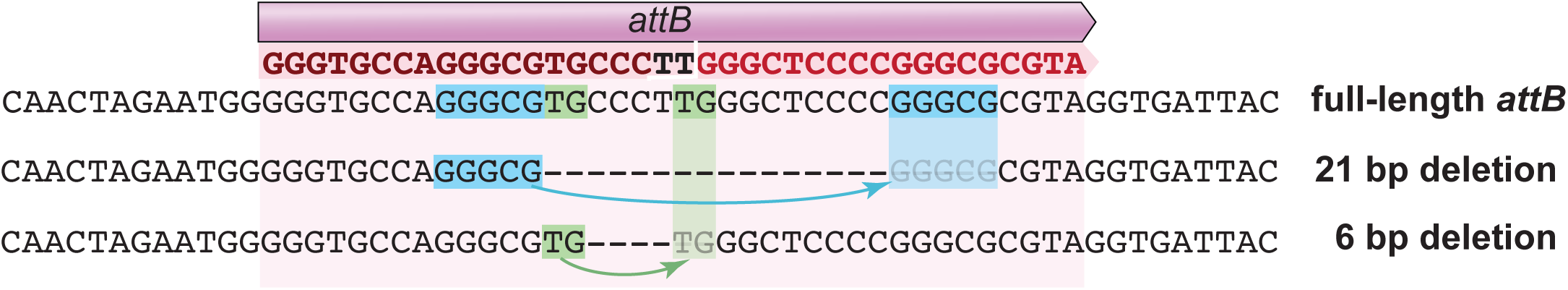
Degraded *attB* from PhiC31 exposure. An *attB* landing pad at the N-terminus of *unc–32* (*attB::unc-32, ox1595*) was allowed proliferate with PhiC31 expressed in the germline for 1-2 months. Propagation of the original strain led to two derived strains with deletions in the *attB* sites. The deletion end points linked short regions of homology (green or blue boxes). It is possible that PhiC31 bound to the *attB* site leads to either DNA damage or replication stress that is repaired by microhomology-mediated end joining. In addition, the high affinity of PhiC31 for the *attB* site (K_d_ 5-20 nM) (McEwan et al., 2011) might interfere with transcription of *unc-32* and provide a selective advantage for the deleted variants.

## Acknowledgements

We thank Mike Nonet for gifts of PhiC31 and Bxb1 expression constructs, in addition to helpful discussions and suggestions, Christian Frøkjaer-Jensen for continuing advice for germline expression. We thank Wayne Davis for reading the manuscript. We thank Matt Schwartz, Christie Wnukowski and Sean Merrill for DNA constructs. EMJ is an investigator of the Howard Hughes Medical Institute. This work was funded by NIH grants R01GM095817 and R01NS03430. MSR was supported by NIH grant F32GM133139.

---

**Table S1.**
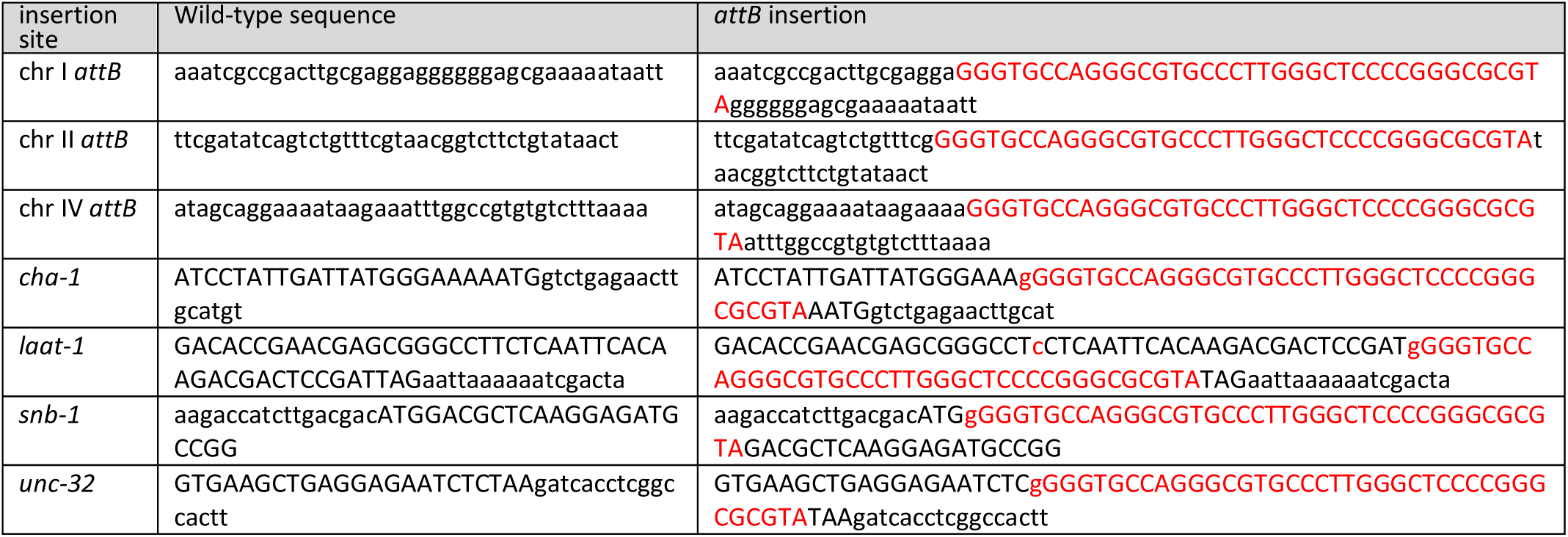

**Table S2.** Strains List.

| attB landing pad strains |  |  |
| --- | --- | --- |
| Strain | Description | Genotype |
| EG10204 | chr II attB landing pad | *oxSi1270[attB] II ; unc-119(ox819) III* |
| EG10250 | chr I attB landing pad | *oxSi1337[attB] I ; unc-119(ox819) III* |
| EG10251 | chrIV attB landing pad | *unc-119(ox819) III ; oxSi1338[attB] IV* |
| EG10463 | laat-1::attB landing pad | *laat-1(ox1141[laat-1::attB]) II ; unc-119(ox819) III* |
| EG10288 | attb::snb-1 landing pad | *unc-119(ox819) III ; snb-1(ox1124[phicattB::snb-1]) V* |
| EG10274 | unc-32::attB landing pad | *unc-32(ox1125[unc-32::phic31attB]) III unc-119(ox819) III* |
| EG10245 | internal cha-1::attB landing pad | *unc-119(ox819) III ; cha-1(ox1298[cha-1::attB]) IV* |
| Excision marker strains |  |  |
| Strain | Description | Genotype |
| EG10035 | attP-sqt-1(e1350) dominant Rol-attB (PhiC31) | *oxSi1430[attP sqt-1(e1350) attB] I* |
| EG10036 | attP-sqt-1(e1350) dominant Rol-attB (Bxb1) | *oxSi1431[Bxb1(attP) sqt-1(e1350) \*jsTi1453] I* |
| EG10040 | loxP-sqt-1(e1350) dominant Rol-loxP | *oxSi1432[loxP sqt-1(e1350) loxP \*jsTi1453] I* |
| EG10043 | rox-sqt-1(e1350) dominant Rol-rox | *oxSi1218[rox sqt-1(e1350) rox \*jsTi1453] I* |
| EG10042 | vox-sqt-1(e1350) dominant Rol-vox | *oxSi1217[vox sqt-1(e1350) vox \*jsTi1453] I* |
| EG10041 | B3RT1-sqt-1(e1350) dominant Rol-B3RT1 | *oxSi1216[B3RT1 sqt-1(e1350) B3RT1 \*jsTi1453] I* |
| EG10388 | B3RT2-sqt-1(e1350) dominant Rol-B3RT2 | *oxSi1239[B3RT2 sqt-1(e1350) B3RT2 \*jsTi1453] I* |
| EG10078 | KDRT-sqt-1(e1350) dominant Rol-KDRT | *oxSi1225[[KDRT::sqt1(e1350)::KDRT \*jsTi1453] I* |
| EG10046 | FRT-sqt-1(e1350) dominant Rol-FRT | *oxSi1433[frt sqt-1(e1350) frt \*jsTi1453] I* |
| Recombinase strains |  |  |
| Strain | Description | Genotype |
| EG10420 | germline expressed Vika | *oxSi1228[loxP Pmex-5::vika::sl2::mNeongreen \*jsSi1579] II* |
| EG10441 | germline expressed B3 | *oxSi1284[loxP Pmex-5::B3::sl2::mNeongreen \*jsSi1579] II* |
| EG10050 | germline expressed Cre | *oxSi1215[Pmex-5::wrmCre-2xNLS::SL2::mNeonGreen::glh-2UTR \*ttTi5605] II* |
| EG10048 | germline expressed FLP | *oxSi1213[loxP Pmex-5-FLP(G5D)::sl2::mNeongreen \*jsSi1579] II* |
| NM5406 | germline expressed PhiC31 | *jsSi1623[loxP Pmex-5::phiC31::SL2::mNeonGreen::glh-2UTR* |
| EG10408 | germline expressed PhiC31 + PATC intron | *oxSi1347[loxP Pmex-5::PhiC31::sl2::mNeongreen \*jsSi1579] II* |
| EG10421 | germline expressed Bxb1 | *oxSi1190[loxP Pmex-5::Bxb1::tbb-2UTR] II* |
| NM5736 | germline expressed Bxb1-nls | *jsSi1834[loxP Pmex-5::Bxb1-NLS::V2A::mNG::glh-2UTR FRT3, \*jsSi1669] IV* |
| EG10378 | germline expressed Vika + tmC27 balancer | *tmC27[unc-75(tmls1239)] I ; oxSi1228[loxP Pmex-5::vika::sl2::mNeongreen \*jsSi1579] II* |
| EG10419 | germline expressed B3 + tmC27 balancer | *tmC27[unc-75(tmls1239)] I ; oxSi1284[loxP Pmex-5::B3::sl2::mNeongreen \*jsSi1579] II* |
| EG10371 | germline expressed Cre + tmC27 balancer | *tmC27[unc-75(tmls1239)] I ; oxSi1215[loxP Pmex-5::wrmCre-2xNLS::SL2::mNeonGreen::glh-2UTR Cbr-unc-119(+)\*ttTi5605] II* |
| EG10370 | germline expressed FLP + tmC27 balancer | *tmC27[unc-75(tmls1239)] I ; oxSi1213[loxP Pmex-5-FLP(G5D)::sl2::mNeongreen \*jsSi1579] II* |
| EG10375 | germline expressed B3 PhiC31 + tmC27 balancer | *tmC27[unc-75(tmls1239)] I ; jsSi1623[loxP Pmex-5::phiC31::SL2::mNeonGreen::glh-2UTR FRT3] IV* |
| EG10372 | germline expressed PhiC31(PATC) + tmC27 balancer | *tmC27[unc-75(tmls1239)] I ; oxSi1347[loxP Pmex-5::PhiC31(PATC)::sl2::mNeongreen \*jsSi1579] II* |
| EG10373 | germline expressed Bxb-1 + tmC27 balancer | *tmC27[unc-75(tmls1239)] I ; oxSi1190[loxP Pmex-5::Bxb1::tbb-2UTR] II* |
| EG10376 | germline expressed Bxb-1-nls + tmC27 balancer | *tmC27[unc-75(tmls1239)] I ; jsSi1834[loxP Pmex-5::Bxb1-NLS::V2A::mNG::glh-2UTR FRT3, \*jsSi1669] IV* |
| PhiT transgene strains |  |  |
| Strain | Description | Genotype |
| EG10379 | chr I dopamine neuron mKate2 (Pdat-1::mKate2(rox)) | *oxSi1378[attL loxP Pdat-1::mKate2::let-858 3'UTR rox::attR \*oxSi1337] I* |
| EG10380 | chr II dopamine neuron mKate2 (Pdat-1::mKate2(rox)) | *oxSi1382[attL loxP Pdat-1::mKate2::let-858 3'UTR rox::attR \*oxSi1270]* |
| EG10381 | chr IV dopamine neuron mKate2 (Pdat-1::mKate2(rox)) | *oxSi1367[attL loxP Pdat-1::mKate2::let-858 3'UTR rox::attR \*oxSi1338] IV* |
| EG10422 | chr II dopamine neuron mKate2 (Pdat-1::mKate2(rox, linear)) | *oxSi1354[attL loxP Pdat-1::mKate2::let-858 3'UTR rox::attR \*ox1270] II* |
| EG10382 | chr IV dopamine neuron mKate2 (Pdat-1::mKate2(B3RT1)) | *oxSi1377[attL loxP Pdat-1::mKate2::let-858 3'UTR B3RT1::attR loxP \*oxSi1338] IV* |
| EG10383 | chr II PVD neuron mKate2 (Pnpr-2::mKate2(rox)) | *oxSi1339[attL loxP Pnpr-2::mKate2::let-858 3'UTR rox::attRP \*oxSi1270]* |
| EG10384 | chr II pan-neuronal mKate2 (Pegl-21::mKate2(rox)) | *oxSi1352[attL loxP Pegl21::mKate2::let-858 3'UTR rox::attR \*ox1270] II* |
| EG10276 | chr II germline expressed PhiC31 + B3 (Pmex-5::PhiC-31::mNeongreen::Pmex-5::B3) | *oxSi1356[attL loxP Pmex-5::B3::let-858 glh-2::mNeongreen::sl2::PhiC31::Pmex-5 \*oxSi1270] II ; unc-119(ox819) III* |

| PhIT tag strains |  |  |
| --- | --- | --- |
| Strain | Description | Genotype |
| EG10424 | mKate2 tagged snb-1 (rox, n-term, injection) | *snb-1(ox1483[attL::mKate2::rox::attR::snb-1 \*ox1124]) V* |
| EG10423 | GFP tagged snb-1 (rox, n-term, injection) | *snb-1(ox1481[attL::GFP::rox::attR::snb-1 \*ox1124]) V* |
| EG10426 | mKate2 tagged unc-32 (rox, c-term, injection) | *unc-32(ox1450[unc-32::attL::mkate2::stop::rox::attR \*ox1125])* |
| EG10425 | GFP tagged unc-32::GFP (rox, c-term, injection) | *unc-32(ox1482[unc-32::attL::GFP::stop::rox::attR \*ox1125]) III* |
| EG10440 | GFP tagged unc-32::GFP (B3RT1, c-term, injection) | *unc-32(ox1512[unc-32::attL::GFP::stop::B3RT1::attR \*ox1125]) III* |
| EG10427 | mKate2 tagged unc-32 (B3RT2, c-term, injection) | *unc-32(ox1452[unc-32::attL::mkate2::stop::b3RT2::attR \*ox1125]) IV* |
| EG10428 | GFP tagged GFP::B3RT2::snb-1 (B3RT2, n-term, injection) | *snb-1(ox1451[attL::GFP::B3RT2::attR::snb-1 \*ox1124]) V* |
| EG10430 | mKate2 tagged snb-1 (B3RT2, n-term, cross) | *snb-1(ox1487[attL::mKate2::B3RT2::attR::snb-1 \*ox1124]) V* |
| EG10429 | GFP tagged snb-1 (B3RT2, n-term, cross) | *snb-1(ox1485[attL::GFP::B3RT2::attR::snb-1 \*ox1124]) V* |
| EG10432 | mKate2 tagged unc-32 (B3RT2, c-term, cross) | *unc-32(ox1488XK_20-23-1_unc-32::attB\*225(1)\*169) III* |
| EG10431 | GFP tagged unc-32 (B3RT2, c-term, cross) | *unc-32(ox1486[unc-32::attL::GFP::stop::b3RT2::attR \*ox1125]) III* |
| EG10312 | AID::GFP tagged cha-1 with neuronal Tir1(579G) (B3RT2, internal, cross) | *cha-1(ox1358[cha-1::attL::aid::gfp::b3rt2::attR \*ox1298]) IV*<br>*oxSi1275[Psnt-1::TIR1(F79G)::F2A::AID::BFP] IV* |
| EG10439 | FLP-on mKate2 tagged snb-1 (rox, n-term, injection) | *unc-119(ox819) III ; snb-1(ox1380[attL::mKate2::FLPon::rox::attR::snb-1 \*ox1124 + loxP]) V* |
| EG10438 | FLP-on Skykan-S tagged laat-1 (rox, c-term, injection) | *laat-1(ox1262[laat-1::attL::FLPon::Skykan-S::rox + loxP]) II*<br>*unc-119(ox819) III* |
| Flp-On strains with tissue specific FLP |  |  |
| Strain | Description | Genotype |
| EG10385 | FLP-on Skykan-S tagged laat-1 + intestinal FLP | *laat-1(ox1262[laat-1::attL::FLPon::Skykan-S::rox::attR + loxP]) II ;*<br>*bqSi508[elt-2p::FLP D5 + unc-119(+)] IV* |
| EG10436 | FLP-on Skykan-S tagged laat-1 + hypodermal FLP | *laat-1(ox1262[laat-1::attL::FLPon::Skykan-S::rox::attR + loxP]) II ;*<br>*bqSi548[dpy-7p::FLP D5 + unc-119(+)] IV* |
| EG10389 | FLP-on mKate2 tagged snb-1 + GABA neuron FLP | *bqSi542[unc-47p::FLP D5 + unc-119(+)] IV ; snb-1(ox1380[attL::mKate2::FLPon::rox::attR::snb-1 + loxP \*ox1124]) V* |
| EG10390 | FLP-on mKate2 tagged snb-1 + pan-neuronal FLP | *oxSi1168[Psnt-1::2xnlS-FLP(G5D) unc-119(-)] II ; snb-1(ox1380[attL::mKate2::FLPon::rox::attR::snb-1 \*ox1124 + loxP]) V* |
| BN528 | Intestinal FLP with heat-shock expression construct | *bqSi294 II ; bqSi508 IV.* |
| BN544 | GABA-neuron FLP with heat-shock expression construct | *bqSi294 II ; bqSi542 IV.* |
| BN550 | hypodermal FLP with heat-shock expression construct | *bqSi294[hsp16.41p::FRT::mCherry::his-58::FRT::GFP::his-58 + unc-119(+)] II ; bqSi548[dpy-7p::FLP D5 + unc-119(+)] IV* |
| EG10038 | pan-neuronal FLP | *oxSi1168[Psnt-1::2xnlS-FLP(G5D) unc-119(-)] II* |
| Extrachromosomal arrays for tagging crosses |  |  |
| Strain | Description | Genotype |
| EG10319 | N-term mkate2 tagging array (B3RT2) | *unc-119(ox819) III ; oxEx2300[attP::mkate2::B3RT2, lssOrange, HisCl channel, him-8 piRNA]* |
| EG10433 | C-term mkate2 tagging array (B3RT2) | *unc-119(ox819) III ; oxEx2301[attP::mkate2::stop::B3RT2, lssOrange, HisCl channel, him-8 piRNA]* |
| EG10434 | N-term GFP tagging array (B3RT2) | *unc-119(ox819) III ; oxEx2302[attP::GFP::B3RT2, lssOrange, HisCl channel, him-8 piRNA]* |
| EG10435 | C-term GFP tagging array (B3RT2) | *unc-119(ox819) III ; oxEx2303[attP::GFP::stop::B3RT2, lssOrange, HisCl channel, him-8 piRNA]* |
| EG10437 | N-term AID::GFP tagging array (B3RT2) | *unc-119(ox819) III ; oxEx2267[attP::AID::GFP::B3RT2, lssOrange, HisCl channel, him-8 piRNA]* |
| RMCE injection strains |  |  |
| Strain | Description | Genotype |
| NM5322 | chr I RMCE landing pad | *jsSi1570 I ; bqSi711 IV.* |
| NM5304 | chr II RMCE landing pad | *jsSi1579[loxP Prpl-28::FRT::GFP::his-58::FRT3] II ; unc-119(ed3) III ;*<br>*bqSi711[Pmex-5::FLP::SL2::mNeonGreen, unc-119(+)] IV* |
| MosSCI injection strains |  |  |
| Strain | Description | Genotype |
| EG6699 | chr II MosSCI landing pad | *ttTi5605 II ; unc-119(ed3) III ; oxEx1578.* |
| EG8078 | chr I universal MosSCI landing pad | *oxTi185 I ; unc-119(ed3) III.* |

